# Ca^2+^-mediated higher-order assembly of b^0,+^AT–rBAT is a key step for system b^0,+^ biogenesis and cystinuria

**DOI:** 10.1101/2021.05.06.443019

**Authors:** Yongchan Lee, Pattama Wiriyasermkul, Satomi Moriyama, Deryck J. Mills, Werner Kühlbrandt, Shushi Nagamori

## Abstract

Cystinuria is a genetic disorder characterized by overexcretion of dibasic amino acids and cystine, which causes recurrent kidney stones and occasionally severe kidney failure. Mutations of the two responsible proteins, rBAT and b^0,+^AT, which comprise system b^0,+^, are linked to type I and non-type I cystinuria respectively and they exhibit distinct phenotypes due to protein trafficking defects or catalytic inactivation. Although recent structural insights into human b^0,+^AT–rBAT suggested a model for transport-inactivating mutations, the mechanisms by which type I mutations trigger trafficking deficiencies are not well understood. Here, using electron cryo-microscopy and biochemistry, we discover that Ca^2+^-mediated higher-order assembly of system b^0,+^ is the key to its trafficking on the cell surface. We show that Ca^2+^ stabilizes the interface between two rBAT molecules to mediate super-dimerization, and this in turn facilitates the N-glycan maturation of system b^0,+^. A common cystinuria mutant T216M and mutations that disrupt the Ca^2+^ site in rBAT cause the loss of higher-order assemblies, resulting in protein trafficking deficiency. Mutations at the super-dimer interface reproduce the mis-trafficking phenotype, demonstrating that super-dimerization is essential for cellular function. Cell-based transport assays confirmed the importance of the Ca^2+^ site and super-dimerization, and additionally suggested which residues are involved in cationic amino acid recognition. Taken together, our results provide the molecular basis of type I cystinuria and serve as a guide to develop new therapeutic strategies against it. More broadly, our findings reveal an unprecedented link between transporter oligomeric assembly and trafficking diseases in general.

## Introduction

Cystinuria is an inherited disorder in which defective amino acid reabsorption in the kidney causes over-excretion of cystine and dibasic amino acids into urine (*1*). Due to the poor solubility of cystine, patients suffer from recurrent kidney stones, which can grow to several centimeters, causing acute pain and kidney failure (*2*). Many patients thus require life-long care, and studies estimate that 1 in 7,000 newborns are affected by the disease (*1*). Common treatments include dietary therapy (*3*), extracorporeal shock wave lithotripsy, management of cystine dilution (*4*) and open surgery, but so far no breakthrough has been made beyond these symptomatic measures due to an incomplete understanding of the disease mechanisms.

Cystinuria is caused by mutations in system b^0,+^, which is a Na^+^-independent cystine/dibasic amino acid exchanger localized at brush border cells in the kidney and intestine. System b^0,+^ is composed of two subunits, a single-transmembrane glycoprotein rBAT (*5*) (also known as D2, NBAT and SLC3A1) and a 12-transmembrane transporter b^0,+^AT (*6*) (also known as BAT1 and SLC7A9), which belong to the heteromeric amino acid transporter (HAT) family (*7*). Although recent structural elucidation of the prototypical HAT transporter, LAT1–CD98hc, has shown how two subunits form a tight heterodimeric assembly (*8, 9*), the low sequence identity of rBAT to CD98hc (28% in the aligned region) and the insertion of a total of ∼100-residues (*10*) have precluded a detailed molecular understanding of b^0,+^AT– rBAT.

Clinical and biochemical studies have shown that mutations of the two subunits are linked to different phenotypes, known as type I cystinuria for rBAT and non-type I for b^0,+^AT (*1*). Whereas most type I mutations cause system b^0,+^ malfunction (*11*), non-type I mutations abolish the transport activity itself (*12*). In addition, while all characterized type I mutations show the disease phenotype only in homozygotes, some non-type I mutations can trigger the disease even in heterozygous individuals (*13*), a phenomenon that currently lacks a clear explanation. Recently, the cryo-EM structures of human b^0,+^AT–rBAT in detergent revealed its architecture and suggested a mechanism of amino acid antiport and its dysfunctions (*14, 15*). However, these structures did not explain why certain type I mutations cause protein trafficking defects.

To better understand the molecular mechanisms of system b^0,+^ biogenesis and cystinuria, we studied the structure and function of ovine system b^0,+^. We determined the electron cryo-microscopy (cryo-EM) structure of ovine b^0,+^AT–rBAT in lipid nanodiscs, which unveils the super-dimeric (b^0,+^AT–rBAT)_2_ complex embedded in a curved lipid bilayer. We identified a Ca^2+^-binding site in the rBAT ectodomain, and demonstrate that this site is essential for higher-order assembly and N-glycan maturation of system b^0,+^. Furthermore, we found that the loss of higher-order assembly is correlated with some type I mutations, revealing a previously unknown link between protein oligomerization and the trafficking disease. These results shed light on the molecular mechanisms of type I cystinuria.

## Results

### Mammalian b^0,+^AT–rBAT complexes form a super-dimer

We purified ovine and murine b^0,+^AT–rBAT complexes, which share 80–88% sequence identities with the human counterparts (Fig. S1) and showed good solution behavior in detergent (Fig. S2a). SDS-PAGE analyses indicated a heterodimer band (∼130 kDa) under non-reducing conditions (Fig. S2b), which separated into two bands for b^0,+^AT and rBAT under reducing conditions, confirming the formation of a disulfide-linked complex. Negative-stain electron microscopy showed two rBAT ectodomains arranged head-to-head, indicative of a higher-order assembly (Fig. 1a). In addition, each ectodomain of rBAT is substantially larger than that of LAT1–CD98hc (Fig. 1b), reflecting the ∼100 amino acid insertion. To verify the functional integrity of the purified complex, we reconstituted ovine b^0,+^AT–rBAT into liposomes (Fig. S2d) and measured amino acid transport activities (Fig. 1c). b^0,+^AT–rBAT showed significant □-[^3^H]Arg uptake when liposomes were loaded with □-Arg. Transport activity was negligible when the liposomes were not loaded with □-Arg, confirming the □-Arg/□-Arg exchange activity (Fig. 1c).

**Figure 1.**
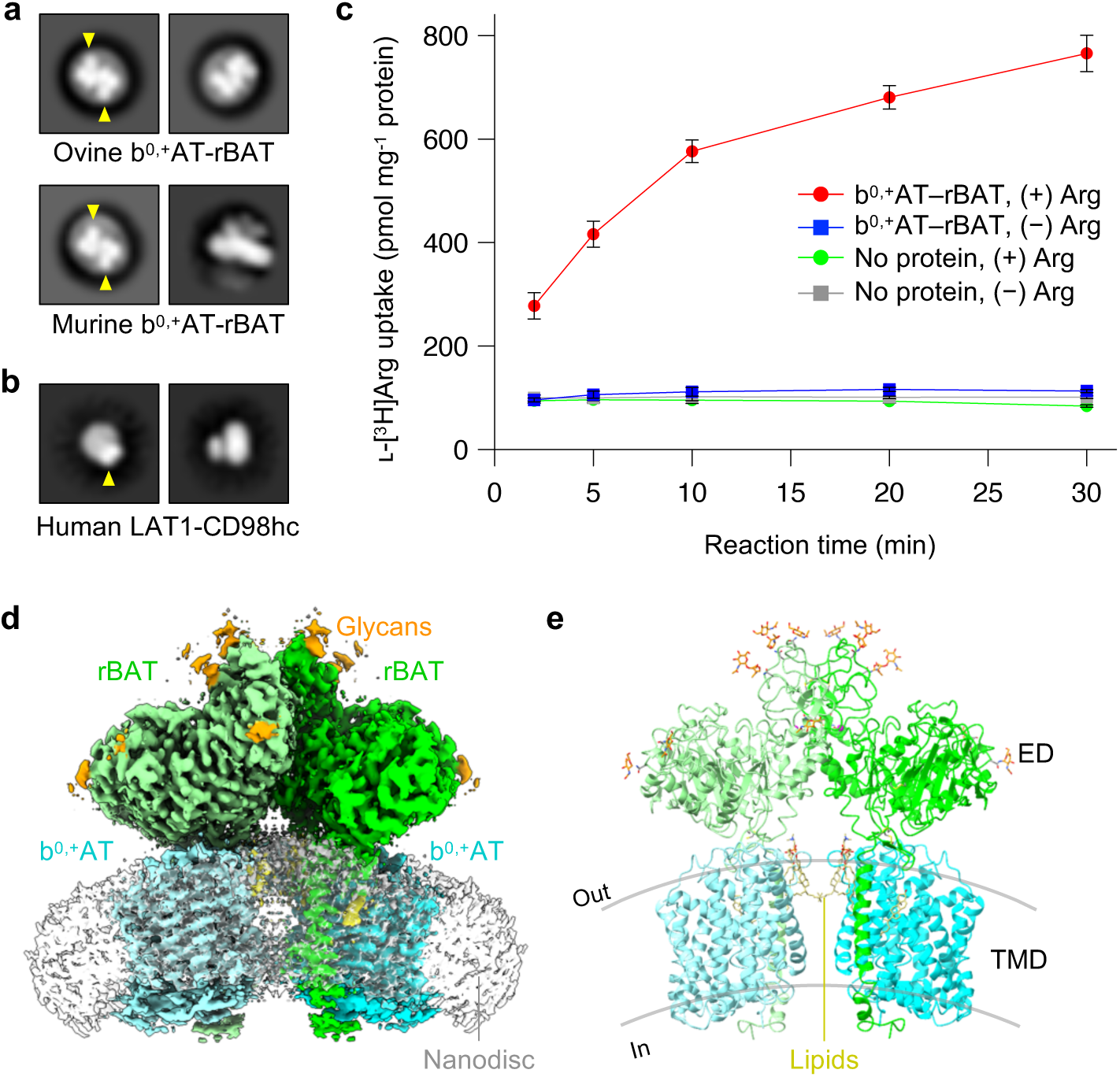
Structural and biochemical characterization of b^0,+^AT–rBAT in a lipid environment. **a)** 2D class averages of ovine and murine b^0,+^AT–rBAT imaged by negative-stain electron microscopy. The rBAT ectodomains are marked by yellow arrows. **b)** 2D class averages of LAT1–CD98hc. The CD98hc ectodomain is marked by a yellow arrow. **c)** Uptake of ʟ-[^3^H]Arg by ovine b^0,+^AT–rBAT reconstituted into proteoliposomes. Proteoliposomes were pre-loaded with either 1 mM or no ʟ-Arg. Control experiments were performed with protein-free liposomes. Values are mean ± s.e.m. n = 3 technical replicates. **d)** Cryo-EM map of the super-dimeric b^0,+^AT–rBAT complex in lipid nanodiscs. Map densities corresponding to different chains and chemical identities are colored as follows: rBAT (green and light green), b^0,+^AT (cyan and light cyan), nanodisc (translucent white), lipids (yellow) and N-linked glycans (orange). **e)** Ribbon diagram of b^0,+^AT–rBAT. The position of the lipid bilayer is based on the lipid-solvent boundaries calculated by the PPM server (https://opm.phar.umich.edu/ppm_server).

Initial single-particle cryo-EM analysis of ovine b^0,+^AT–rBAT complex in detergent yielded a map at an overall resolution of 3.9 Å (Fig. S3a). After rigorous optimization, we reconstituted the complex into lipid nanodiscs consisting of MSP1E3D1, phospholipids and cholesterol (Fig. S2c, see Methods for detail), which yielded a better map at 2.9 Å global resolution (Fig. S3b–f). To further improve the map quality, we performed the multi-body refinement and focused refinement (*16*), improving the map resolution to 2.6 Å for the extracellular domain and 3.0 Å for the individual heterodimer (Fig. S3e,f). These maps allowed us to build an atomic model of ovine b^0,+^AT–rBAT (Figs. S4a–e and S5).

### Overall structure of the ovine b^0,+^AT–rBAT complex

The overall structure of the ovine system b^0,+^ shows two copies of b^0,+^AT and rBAT, arranged as a super-dimer of the heterodimer (Fig. 1d–e). This architecture is essentially identical to the recent structures of human b^0,+^AT–rBAT (*14*), confirming a conserved higher-order assembly (Fig. S6b). rBAT has three domains, namely the ectodomain, a single transmembrane helix (TM1’) and the cytoplasmic N-terminal helix (NH) (Fig. 2a–c), in addition to 62 unresolved disordered residues at the N-terminus. The ectodomain displays a glucosidase-like fold (*10, 17*), which consists of domains A, B and C (Fig. 2b). The cryo-EM map also resolved a Ca^2+^ ion and six N-linked glycans for each rBAT molecule, as discussed later in detail.

**Figure 2.**
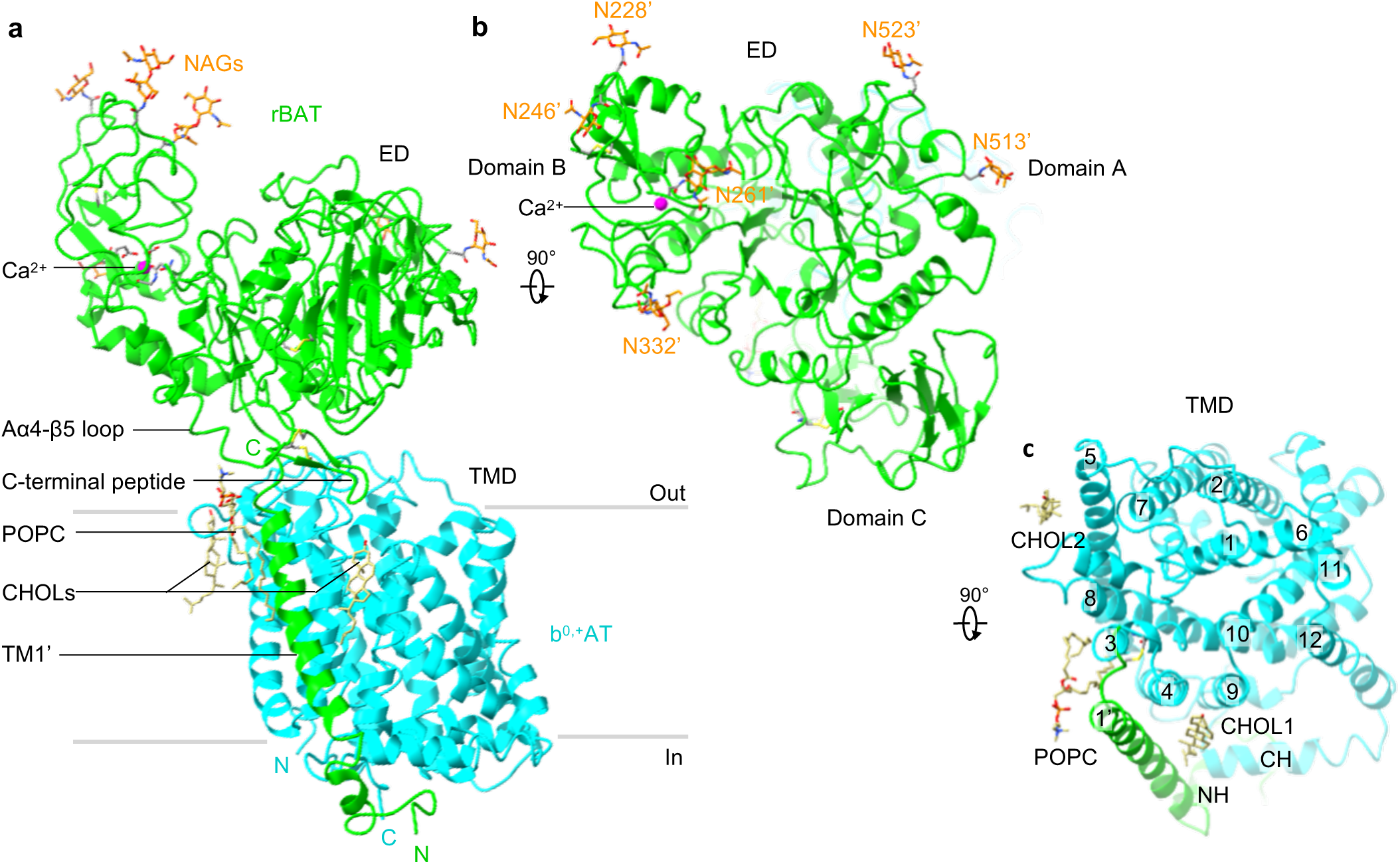
Structure of the ovine b^0,+^AT–rBAT heterodimer. **a)** Detailed depiction of the b^0,+^AT–rBAT subcomplex, derived from 3D multi-body refinement (see Methods). Note that the complex is re-oriented from Fig. 1e to the membrane normal. CHOL, cholesterol; POPC, palmitoyl oleoyl phosphatidyl choline; TMD; transmembrane domain; ED, ectodomain; NAG, N-acetylglucosamine. **b)** The rBAT ectodomain. The Ca^2+^ ion, N-linked glycans and individual subdomains are labeled. **c)** b^0,+^AT complexed with rBAT TM1’. Lipids and individual TMs are labeled. CH, C-terminal helix; NH, N-terminal helix.

b^0,+^AT shares structural and sequence homology with the APC superfamily and shows a typical LeuT-fold (Fig. 2c), consisting of 12 transmembrane helices (TM1–12) that contain 5 + 5 TM inverted repeats (*18*). Other structural features involve two helices in the extracellular loop 4 (EL4a and EL4b), one helix in intracellular loop 1 (IL1) and the cytoplasmic C-terminal helix (CH) running parallel to the membrane (Fig. S5). Notably, the cryo-EM map resolved numerous lipids, two of which were assigned to cholesterol and one to phospholipid (Fig. 2a, c and Fig. S4d, e). Furthermore, the lipid nanodisc enclosing the TMD is bent by about ∼30 degrees (Fig. 1c), which is reminiscent of the highly-curved membranes of the brush border microvilli, where system b^0,+^ resides (*19*).

### The b^0,+^AT–rBAT interface

Single-particle analysis indicated that the two b^0,+^AT–rBAT subcomplexes are linked together with substantial flexibility, which was also evident in the human complex that showed a blurred TMD map in the consensus 3D refinement (*14, 15*). To account for this flexibility, we performed multi-body refinement (*16*), taking individual heterodimers as rigid bodies moving relative to each other. This analysis not only revealed the swinging motions of b^0,+^AT relative to the rigid rBAT core (Movie S1), but also yielded an improved map for the individual b^0,+^AT–rBAT subcomplex at 3.0-Å nominal resolution (Fig. S3c,f). This new map covered the intact b^0,+^AT–rBAT heterodimer, which was not the case in the previous studies (*14, 15*). It therefore allowed us to analyze the b^0,+^AT–rBAT interface in greater detail (Fig. 3).

**Figure 3.**
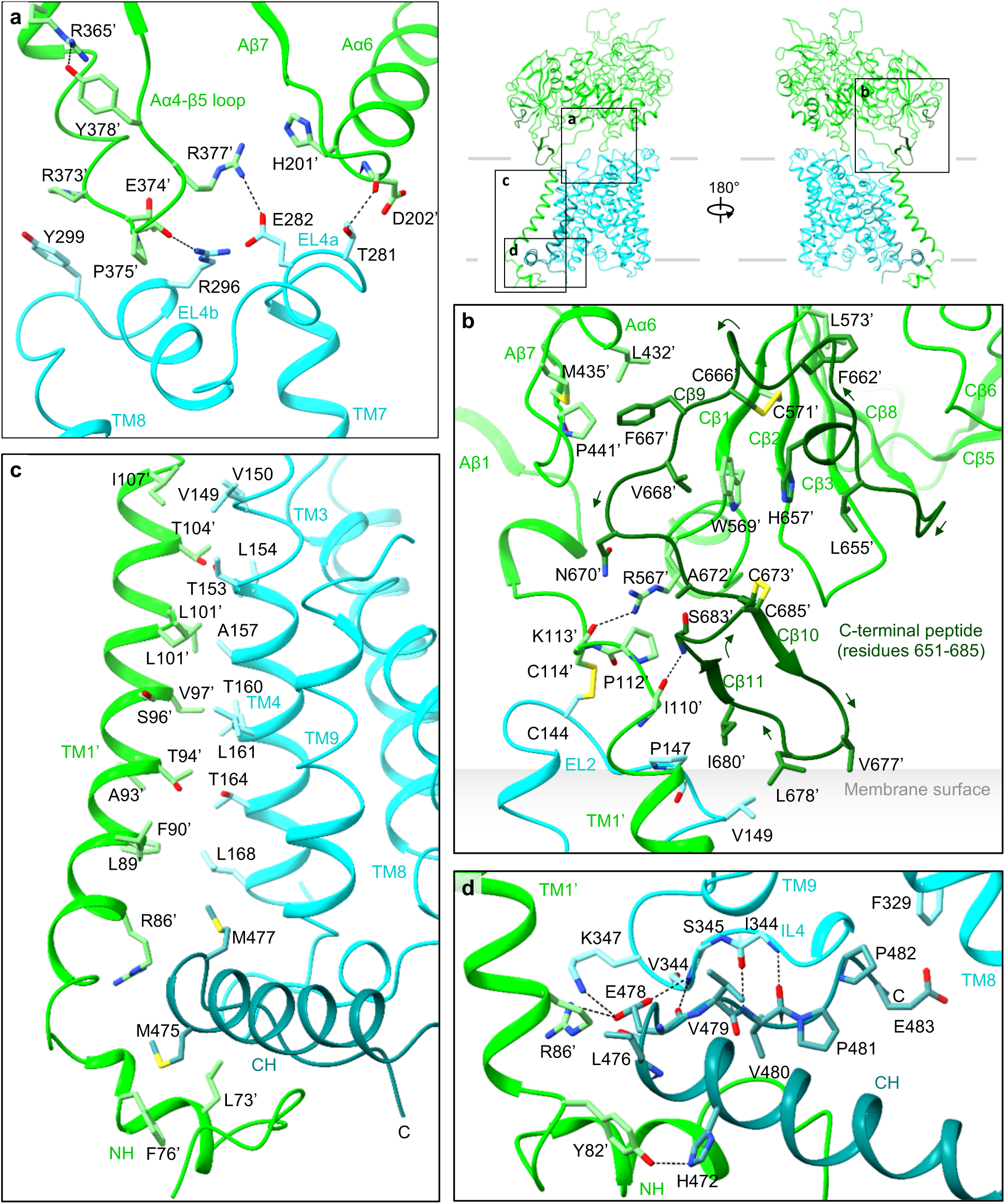
The interface analysis of b^0,+^AT–rBAT. **a)** Extracellular interactions formed by the Aα4-β5 loop, Aα6, EL4a and EL4b. Key interacting residues are shown, with hydrophilic interactions depicted by dotted lines. Arg365’ stabilizes the Aα4-β5 loop through interaction with Tyr378’. The locations of panels **a–d** are indicated on the right. **b)** C-terminal peptide of rBAT. The peptide is dark green for better visibility. Leu678’ contacts the membrane surface, as calculated by the PPM server (https://opm.phar.umich.edu/ppm_server). **c)** Intramembrane interactions. TM1’ and TM4 have numerous hydrophobic residues that pack against each other. The C-terminal helix (CH) of b^0,+^AT is dark cyan for better visibility. **d)** CH and ‘Val-Pro-Pro’ motif of b^0,+^AT, forming intracellular interactions.

On the extracellular side, EL2 of b^0,+^AT is disulfide-bonded to the linker that connects TM1’ to the ectodomain of rBAT (Fig. 3b). EL4a and EL4b form salt bridges and hydrogen bonds to the Aα4-β5 loop and Aα6 (Fig. 3a). Notably, the Aα4-β5 loop is stabilized by Arg365’ (Fig. 3a), a residue associated with a prevalent cystinuria mutation R365W (*1*), suggesting that this substitution disrupts the b^0,+^AT–rBAT extracellular interface. The extracellular interaction is further strengthened by the C-terminal peptide of rBAT (residues 651–685), which is absent in CD98hc (Fig. 3b) (*8, 9*). The first half of the peptide wraps around Cβ1–3 to form the ninth β-strand (Cβ9), which connects domain A and C (Fig. 3b). The second half folds into a cyclic β–hairpin, which interacts with b AT and touches the lipid bilayer (Fig. 3b). Leu678’, which is the site of a known cystinuria mutation L678P (*5*), is located at the tip of this hairpin, suggesting a role in complex stabilization or membrane anchoring.

Within the membrane, TM1’ and TM4 form extensive helix-helix packing interactions (Fig. 3c). We identified one cholesterol molecule bound tightly at this interface (Figs. 2c and S4a,d,e), which is in an equivalent position to that seen in LAT1–CD98hc (*9*), suggesting a conserved cholesterol binding site in the HATs. Located near the cholesterol binding site is Leu89’ (Fig. S4a), which, when mutated to L89P, causes inefficient assembly with b^0,+^AT and cystinuria (*11*), highlighting its importance for heteromeric interactions. On the cytoplasmic side, the N-terminal helix (NH) of rBAT interacts with the C-terminal helix (CH) of b^0,+^AT (Fig. 3d). The CH is unwound at Glu478. The subsequent residues extend into the cleft between NH and IL4 and are partially exposed to the cytoplasm (Fig. 3d). These residues correspond to the ^480^Val-Pro-Pro^482^ motif, which has been suggested to interact with currently unknown cytoplasmic factors to regulate ER–Golgi trafficking (*20*). Pro482, a residue frequently mutated in P482L that gives rise to a Japanese form of cystinuria (*12*), forms van-der-Waals interaction with IL4 (Fig. 3d and Fig. S4b), highlighting its structural importance.

We previously noted that the heterodimerization interface might differ between HAT subgroups that associate with either rBAT or CD98hc (*9*). To investigate the difference, we superimposed the structure of b^0,+^AT–rBAT onto LAT1–CD98hc using the TMDs, which showed that two ectodomains are displaced by ∼40 Å to create distinct interfaces (Fig. S6a). Moreover, surface electrostatic potential calculations show that the rBAT ectodomain is mostly negatively charged (Fig. S6c), whereas CD98hc is positively charged (*9*), indicating different electrostatic interactions. In contrast to the distinct extracellular interactions, the transmembrane and cytoplasmic interactions are similar (Fig. S6a). Given the low sequence conservation of TM1’, this observation is remarkable and could suggest strong evolutionary pressure acting on the intramembrane interactions of HATs.

### rBAT subdomains, disulfide bonds and glycosylation

The domain B is unique to rBAT (not present in CD98hc) and comprised of two insertion loops within domain A (Fig. S5). For simplicity, we refer to the first loop as domain B-I (residues 213–289) and the second loop as domain B-II (residues 318–355) (Fig. 4a,b). Domain B-I has three β-strands and one α-helix, and contacts domain A through a cluster of hydrophobic residues (Fig. 4d). Thr216’ is located at the core of van-der-Waals interactions (Fig. 4d), suggesting how a common cystinuria mutation T216M (*1*) would destabilize the domain A–B interface. Likewise, the domain A–C interface has Met467’, which connects domains A and C through hydrophobic interactions (Fig. 4b,e), suggesting how M467T (*1*) would destabilize this interface. Domain B-II interacts with the adjacent rBAT molecule and thereby contributes to the formation of higher-order assemblies (Fig. 4b). The interaction involves six salt bridges, one π-cation-π stacking and several hydrophobic interactions (Fig. 4c), which together give rise to the strong higher-order assembly.

**Figure 4.**
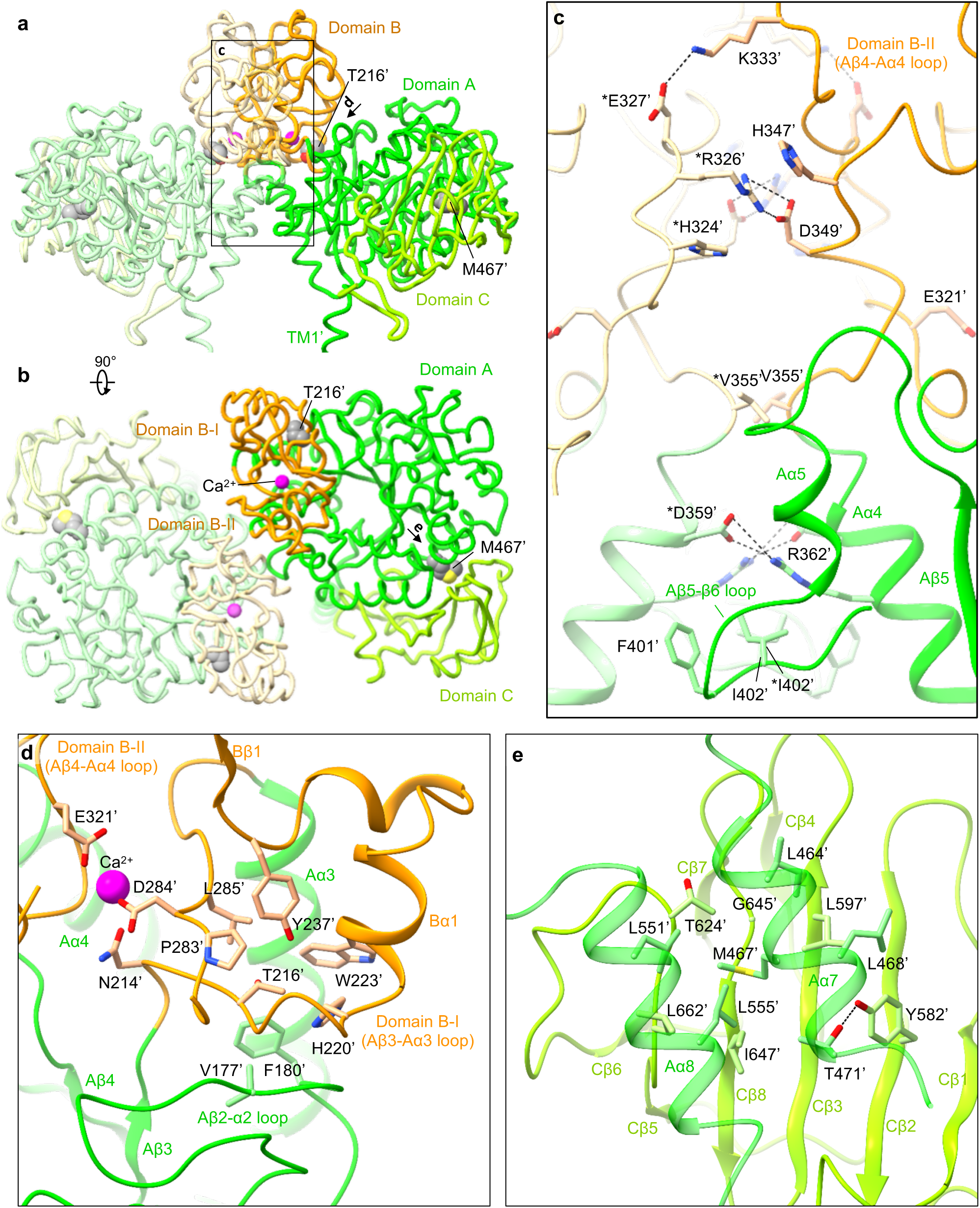
The rBAT ectodomain. **a)** Structure of the ectodomain homo-dimer. The three subdomains are A (green), B (orange) and C (light geren). Common cystinuria-related residues Thr216’ and Met467’ and the Ca^2+^ ions are shown as spheres. **b)** Extracellular view of the rBAT ectodomain. **c)** Zoom-up of super-dimer interface. Salt bridges, π-cation-π stacking and van-der-Waals interactions are indicated. Residues from the adjacent protomer are marked with asterisks (*). **d)** Interaction network around Thr216’ and Ca^2+^ **e)** Interaction network around Met467’.

rBAT has three internal disulfide-bonds (Fig. S5), which are formed during b^0,+^AT-dependent oxidative folding (*21, 22*). The first of these is the functionally most important (*21*) and is formed between Cys242’ and Cys273’, stabilizing domain B (Fig. S7a). The second bond is formed between Cys571’–Cys666’ in domain C, stabilizing the beta-sheet (Fig. S7a). The third bond is formed between Cys673’ and the very C-terminus of rBAT, Cys685’, resulting in a cyclic C-terminal β-hairpin (Figs. 3b). Such a C-terminal disulfide bond is rarely seen in protein structures, highlighting the unique nature of the rBAT ectodomain.

Ovine rBAT has six N-linked glycans, three of which are shared in human (Fig. S7b). Human rBAT has an additional glycan on Asn575’, which is critical for rBAT maturation (*23*). Asn575’ in ovine rBAT is not glycosylated, due to the replacement of Ser577’ to Asn577’ (Fig. S7b), which disrupts the third letter of the glycosylation motif NX(T/S). Structural comparison shows that the non-glycosylated Asn575’ of ovine rBAT adopts a typical β sheet conformation, with its side-chain interacting with domain A (Fig. S7c). By contrast, the glycosylated Asn575’ of human rBAT is flipped out of the β sheet and is exposed to the solvent, with its glycan moiety interacting with domain A (Fig. S7c). This structural difference is caused by the bulky amino acids Tyr425’ and Tyr579’ pushing away Asn575’ in human rBAT, which are replaced by smaller amino acids Ser425’ and Ser579’ in ovine rBAT. Thus, human rBAT has an additional glycan for stabilizing the domain A–C interface, which explains why disrupting this glycan causes premature degradation of system b^0,+^ in humans (*23*).

### Ca^2+^ binding site

The cryo-EM map of rBAT revealed a strong spherical density in the middle of domain B (Fig. 5a). This density is surrounded by the acidic side chains of Asn214’, Asp284’ and Glu321’ and the main chain carbonyl groups of Tyr318’ and Leu319’, indicative of a metal cation (Fig. 5b). Based on the coordination geometry and the sequence homology with known Ca^2+^-binding sites of amylases (*24*), we assigned this density to a bound Ca^2+^ ion. As we did not supply any Ca^2+^ during purification, the bound ion has most likely been acquired during protein expression. This density was also observed in the recent structures of human rBAT and assigned as Ca^2+^, supporting a conserved binding site (*14, 15*).

**Figure 5.**
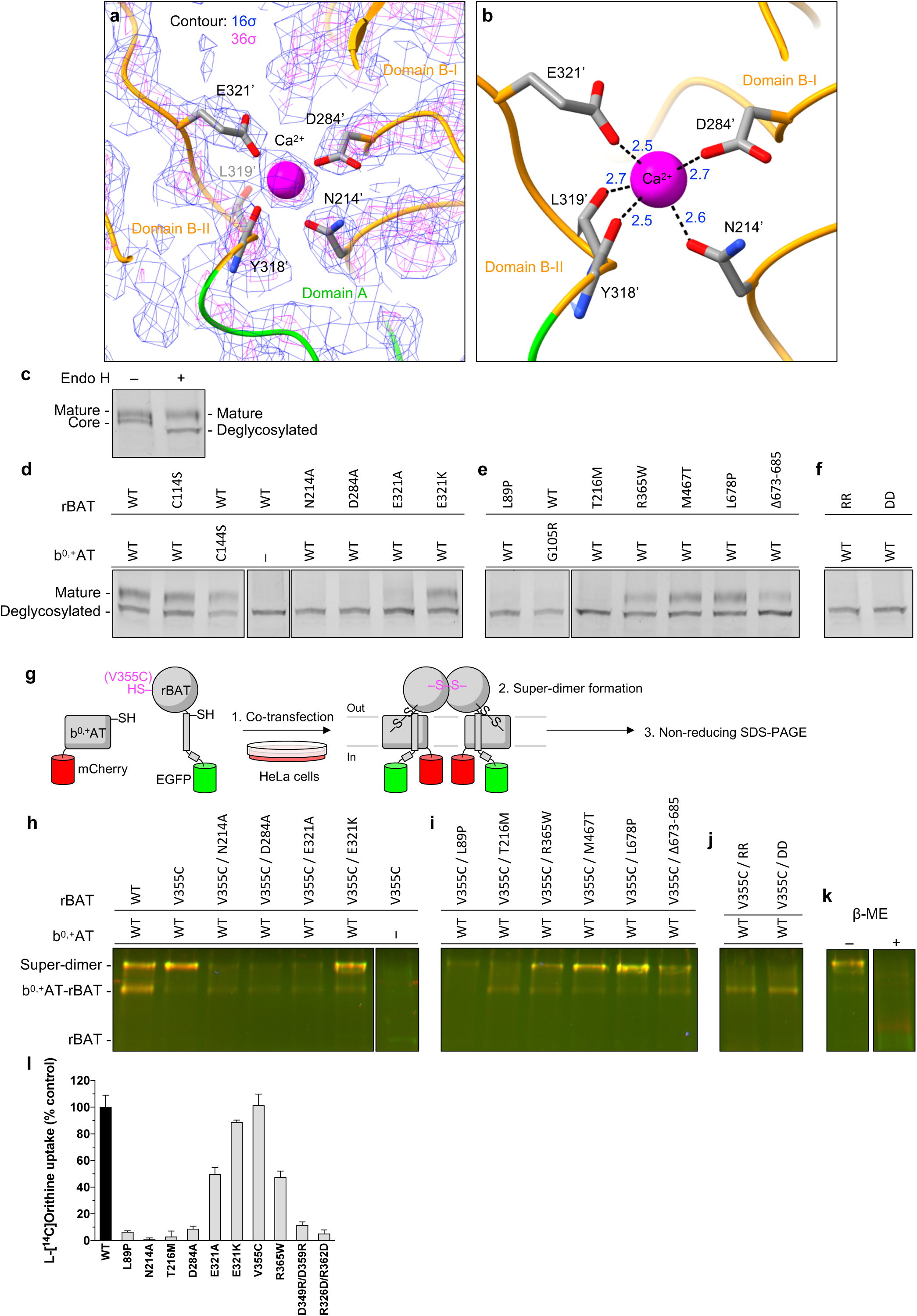
The Ca^2+^ binding site. **a)** Cryo-EM map of the Ca^2+^ binding site at 2.6 Å resolution, contoured at 16 σ (blue) and 36 σ (magenta). **b)** Close-up view of the Ca^2+^ site. The coordination distances in Å are labeled. Note that unmodelled water molecules may contribute to the full Ca^2+^ coordination. **c)** Endo H assay to evaluate the N-glycan maturation. Shown is the fluorescence signal from EGFP fused to rBAT, detected in the SDS-PAGE gel. The wild-type rBAT yields a mature, higher molecular-weight form that resists Endo H treatment (upper band), in addition to the core-glycosylated form (lower band, left lane), which gets de-glycosylated by the Endo H treatment (lowest band, right lane). **d)** Endo H assay for the Ca^2+^ binding site. Lanes 1–3 are positive controls, where WT or disulfide-less mutants (rBAT C114S or b^0,+^AT C144S) were tested for Endo H sensitivity and showed normal rBAT maturation. Lane 4 is a negative control, where rBAT in the absence of b^0,+^AT lost the mature band. Lanes 5– 8 are for Ca^2+^-binding site mutants, where Ala substitutions of the Ca^2+^-coordinating residues abolished the rBAT maturation, whereas E321K restored it nearly to the level of WT. **e)** Endo H assay for major cystinuria mutants (locations shown in **Figs. 3a,b,c and 4d,e**). The mutants L89P, G105R (b^0,+^AT) and T216M showed an almost complete loss of rBAT maturation, whereas R365W, M467T and L678P retained the mature rBAT band to some degrees. The deletion of the C-terminal peptide (Fig. 3b) showed severer effect than a point mutation L678P on this peptide. **f)** Endo H assay for super-dimerization mutants (locations shown in Fig. 4c). The two double mutations D349R/D359R (labeled RR) and R362D/R326D (labeled DD) completely abolished maturation. **g)** The workflow of site-specific cross-linking assay for detecting super-dimers. V355C introduces a pair of cysteines at the rBAT–rBAT homomeric interface (locations shown in Fig. 4c). When b^0,+^AT–rBAT forms a super-dimer, a pair of Cys355’ residues form a disulfide bond, which can be detected as higher-molecular weight species in oxidizing SDS-PAGE. Also see **Fig. S9a–c** for larger gels. **h)** Cross-linking assay for Ca^2+^ site mutants. Overlay of two fluorescence channels imaging non-reducing SDS-PAGE. Green is for EGFP (488 nm), and red is for mCherry (546 nm). Yellow thus represents the b^0,+^AT–rBAT heterocomplexes (monomeric or super-dimreic). Note that wild-type rBAT mutations show some super-dimers even without the V355C mutation, indicating a strong assembly. **i)** Cross-linking assay for selected cystinuria mutants. **j)** Cross-linking assay for super-dimer interface mutants. **k)** Control experiments for cross-linking assay. Addition of β-mercaptoethanol dissociates b^0,+^AT and rBAT into two separate bands. **l)** Uptake of ʟ-[^14^C]-Orn by proteoliposomes reconstituted with b^0,+^AT–rBAT or its mutants. Net uptake was normalized to WT set as 100%. Values are mean ± s.e.m. n = 4 technical replicates.

To gain insight into the Ca^2+^ binding site, we compared the rBAT ectodomain with homologous glucosidases with conserved Ca^2+^ binding sites. A homology search revealed close structural similarity to an α-amylase from *Anoxybacillus* species, known as TASKA (*24*), and indicated that the observed Ca^2+^ site in rBAT corresponds to ‘site 1’ of TASKA, which stabilizes the protruding loop of domain B (*24*). In rBAT, the Ca^2+^ ion bridges domains B-I and B-II to reinforce the super-dimer interface (Fig. 4c,d). In addition, electrostatic potential calculations of rBAT with or without Ca^2+^ showed that Ca^2+^ neutralizes the super-dimer interface to facilitate the homomeric interaction (Fig. S8a,b). Although sequence homology suggested the presence of another Ca^2+^ binding site, known as ‘site 2’ in TASKA (Fig. S7d), this site is occupied by a weaker peak in our cryo-EM map, indicating no or sub-stoichiometric binding (Fig. S7e). Although Ca^2+^ ions regulate the enzymatic activity of amylases (*25, 26*), there is no evidence for any enzymatic activity of rBAT. Therefore, we explored a different physiological role of Ca^2+^.

### The role of Ca^2+^ in system b^0,+^ maturation

Many ER-resident chaperones and post-translational modification machineries, such as calreticulin (*27, 28*), BiP/GRP78 (*29, 30*) and the protein disulfide isomerase (*31*), are known to bind Ca^2+^ as a regulatory element (*32*). Therefore, we hypothesized that the bound Ca^2+^ in rBAT may be involved in the system b^0,+^ biogenesis in the ER. To test this hypothesis, we investigated the N-glycan maturation of system b^0,+^ using the Endo H assay previously established (*11*). We modified the assay by fusing GFP to rBAT and mCherry to b^0,+^AT, to enable fluorescence-based band detection in SDS-PAGE (Fig. S9a,b). As a positive control, we co-expressed the fluorescence-tagged wild-type rBAT and b^0,+^AT, which yielded an Endo H-resistant rBAT band, demonstrating that wild-type rBAT underwent glycan maturation (Fig. 5c; lanes 1 and 2). By contrast, rBAT alone did not mature in the absence of b^0,+^AT (Fig. 5d; lane 4), confirming the requirement of b^0,+^AT for rBAT maturation (*11*). Using this assay, we investigated the effects of the Ca^2+^-binding site mutations, N214A, D284A and E321A, on glycan maturation. Intriguingly, all three mutants showed significantly decreased maturation (Fig.7d; lanes 5–7), demonstrating that Ca^2+^ binding is important for protein maturation. Cellular protein localization confirmed that these mutants cause impaired cell-surface trafficking (Fig. S11).

Given that Ca^2+^ introduces a positive charge to stabilize the cluster of negatively-charged residues, we next asked if adding a positive charge can complement these mutants. To this end, we prepared an E321K mutant, which would mimic the positive charge of Ca^2+^ to compensate for its loss. Indeed, the E321K mutant restored the maturation (Fig. 5d; lane 8) and plasma membrane localization (Fig. S11b), confirming the importance of a positive charge at this position for rBAT maturation. Notably, some amylases naturally have Lys at this position (e.g. Lys206 in a GH13 amylase; PDB 5ZCC) (*33*), suggesting a conserved yet substitutable role of Ca^2+^.

### Domain B and cystinuria

We next asked if any of the known cystinuria mutations exhibit correlated phenotypes to the Ca^2+^ site mutants. To test this, we investigated representative cystinuria mutants from each subdomain, namely L89P (TM1’), T216M (domain B), R365W (the domain A-TMD interface), M467T (the domain A–C interface), L678P (domain C) and b^0,+^AT G105R (TMD) (*1, 34*), in addition to the rBAT C-terminal deletion mutant Δ673-685. Consistent with previous studies (*11, 35*), our Endo H assays showed that L89P on TM1’ abolished glycan maturation (Fig. 5e, lane 1), and G105R resulted in improper b^0,+^AT folding to stop rBAT maturation (Fig. 5e; lane 2), supporting a critical importance of intramembrane interactions for glycan maturation. Interestingly, the mutant T216M in domain B resulted in a complete loss of maturation (Fig. 5e; lane 3), whereas R365W, M467T and L678P in domain A or C showed no or only a slight decrease in maturation (Fig. 5e; lanes 4–6), suggesting that domain B plays a critical role for glycan maturation of system b^0,+^. Deletion of the C-terminal peptide (Δ673-685) resulted in a small decrease, supporting that this peptide is also important for the system b^0,+^ maturation (Fig. 5e: lane 7).

### Super-dimerization is crucial for system b^0,+^ maturation

Given that Ca^2+^ and domain B are both structurally involved in super-dimer formation (Fig. 4c), we reasoned that mutations in domain B or the Ca^2+^ site first affect the super-dimerization and thereby impair protein maturation. To test this assumption, we evaluated super-dimer formation of the mutants using a site-specific disulfide cross-linking assay (Figs. 5g–k and S10). As a site for introducing cysteine, we chose Val355’, which interacts with *Val355’ of the adjacent protomer, with a Cα–Cα distance of ∼4.4 Å (Fig. 4c), suitable for spontaneous disulfide-bond formation. Indeed, when co-expressed with b^0,+^AT in HeLa cells, rBAT V355C showed the cross-linked super-dimer for nearly 100% under non-reducing SDS-PAGE (Fig. 5h; lanes 1–2), confirming site-specific cross-linking. This cross-linking did not occur without b^0,+^AT, indicating that the rBAT–rBAT association requires b^0,+^AT (Fig. 5h; lane 7). To investigate the effect of Ca^2+^ on super-dimer formation, we introduced the mutants N214A, D284A or E321A into this V355C background. In all of them, super-dimer formation was reduced (Fig. 5h; lanes 3–5), revealing the importance of Ca^2+^ for super-dimer formation. The cation-compensating mutation, E321K, restored the super-dimer (Fig. 5h; lane 6), supporting a role of the positive charge for the super-dimer formation.

We next asked if any of the known cystinuria mutations would prevent super-dimerization. Intriguingly, the T216M mutation in domain B mutant showed an almost complete loss of the super-dimer (Fig. 5i; lane 2). By contrast, the R365W, M467T, L678P and Δ673-685 mutations, which are all located in domain A and C, retained super-dimers (Fig. S5i; lanes 3–6). Together with the above Endo H assays, these results suggest that the T216M mutation in domain B disrupts the higher-order assembly and prevents the N-glycan maturation, and thereby leads to cystinuria. Unlike T216M, the L89P mutant formed super-dimers (Fig. 5i; lane 1) but did not mature (Fig. 5e; lane 1), suggesting a different mechanism. L89P may cause protein instability by affecting the b^0,+^AT–rBAT assembly via a conserved cholesterol-binding site (Fig. S4a).

To confirm that super-dimerization *per se* is important for protein maturation, we directly mutated the residues at the super-dimer interface. Two double mutants, D349R/D359R and R362D/R326D, disrupt two critical salt bridges at the rBAT– rBAT interface (Fig. 4c) and would thus abolish the super-dimer. Indeed, when tested with cross-linking assay, both V355C/D349R/D359R and V355C/R362D/R326D did not form super-dimers while retaining the hetero-dimeric assembly (Fig. 5k; lanes 1– 2), validating our mutant design. We then tested these mutants with the Endo H assay, which showed that both mutants did not undergo glycan maturation (Fig. 5f; lanes 1–2). These results confirm our hypothesis that the super-dimerization is the key for rBAT maturation.

To confirm these findings in cellular contexts, we performed amino acid transport assays. We used wild-type b^0,+^AT and rBAT without fluorescent tags and the HEK293 cell line that has the full glycosylation machinery, to reliably evaluate the effects of N-glycan deficient mutations. We set up the assay for the transport of Orn, a physiological substrate of system b^0,+^, and validated the setup with known cystinuria mutants. Consistent with previous observations (*11*), wild-type cells showed significant Orn uptake (Fig. 5l), whereas the three tested mutants (L89P, T216M and R365W) showed no or reduced activities in our assays (Fig. 5l), validating our assay setup. We then tested the mutants of the Ca^2+^ binding site, N214A, D284A and E321A, which showed significantly reduced uptake (Fig. 5l), indicating cellular effects of these mutants. By contrast, the cation-compensating mutant E321K restored activity close to 100%, confirming the functional role of Ca^2+^. Furthermore, the V355C mutant, which was used for the cross-linking assay, showed negligible effect on transport activity, validating our assay. Finally, the super-dimerization mutants (D349R/D359R and R362D/R326D) showed an almost complete loss of activity (Fig. 5l), confirming that the super-dimerization is critical for cellular functions. Altogether, these results revealed an unexpected role of Ca^2+^ for the formation of higher-order assemblies, maturation and cellular function of system b^0,+^.

### Substrate specificity of b^0,+^AT

As in other LeuT-fold transporters, a putative substrate binding site of b^0,+^AT is formed between the hash and bundle domains (*36, 37*) (Fig. 6a,b). At this site, the two broken helices TM1 and TM6 expose the main-chain amino and carbonyl groups to the cytoplasmic solvent (Fig. 6c,d), forming a substrate backbone recognition site. TM6 features Asp233 (Fig. 6d), which renders the site negatively charged, suggesting a role in recognizing positively-charged substrates.

**Figure 6.**
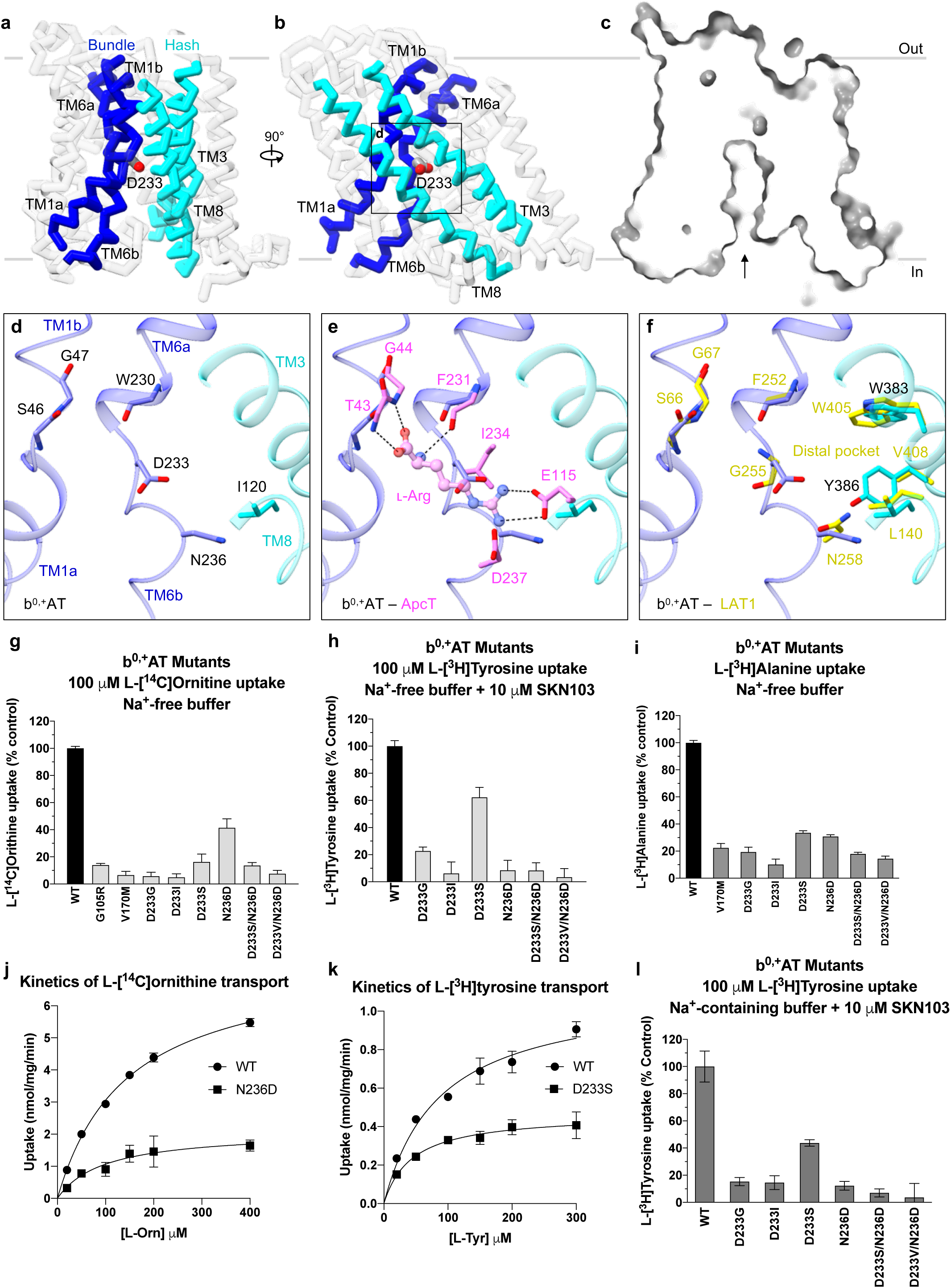
Putative substrate-binding site of b^0,+^AT and transport assays. **a,b)** Cα trace representation of b^0,+^AT, highlighting the TMs involved in substrate translocation. The hash domain is colored cyan and the bundle domain is blue. The Asp233 is shown as stick model. **c)** Cut-away surface representation of b^0,+^AT from the same view as Fig. 6a. The inward-facing cavity is indicated by an arrow. **d)** Close-up view of putative substrate-binding site. **e)** Comparison of b^0,+^AT with ApcT. Residues of ApcT are shown in pink. **f)** Comparison of b^0,+^AT with LAT1. Residues of LAT1 are shown in yellow. **g)** Uptake of ʟ-[^14^C]Orn by b^0,+^AT mutants in HeLa cells expressing b^0,+^AT–rBAT or its mutants. Net uptake was normalized to WT set as 100%. Values are mean ± s.e.m. n = 4 technical replicates. **h)** Uptake of ʟ-[^3^H]Tyr by b^0,+^AT mutants. Values are mean ± s.e.m. n = 4 technical replicates. **i)** Uptake of ʟ-[^3^H]Ala by b^0,+^AT mutants. Values are mean ± s.e.m. n = 4 technical replicates. **j)** Concentration-dependent uptake of ʟ-[^14^C]-Orn by b^0,+^AT WT and N236D. Km and Vmax of ʟ-[^14^C]-Orn transport by b^0,+^AT WT are 145 μM and 7.5 nmol/mg prot./min, respectively, those of N236D are 97 μM and 2.1 nmol/mg prot./min, respectively. **k)** Concentration-dependent uptake of ʟ-[^3^H]-Tyr by b^0,+^AT WT and D233S. Km and Vmax of ʟ-[^3^H]-Tyr transport by b^0,+^AT WT are 87 μM and 1.1 nmol/mg prot./min, respectively, and those of N233S are 44 μM and 0.5 nmol/mg prot./min, respectively. **l)** Uptake of ʟ-[^3^H]Tyr by b^0,+^AT mutants in the presence of external Na^+^. Values are mean ± s.e.m. n = 4 technical replicates.

To gain insight into possible substrate recognition mechanisms, we compared the structure of b^0,+^AT with GkApcT (*38*), a bacterial homolog of the cationic amino acid transporters (CATs; SLC7A1–4) (*39*). In the Arg-bound structure of GkApcT (M321S) (*38*), the substrate backbone is recognized by the main chains of TM1 and TM6 and the substrate guanidium group is recognized by Glu115 on TM8 (Fig. 6e). This Glu115 is not conserved in b^0,+^AT, but instead Asp233 would be located adjacent to the guanidium group (Fig. 6e), supporting its involvement in the positive charge recognition. GkApcT has another acidic residue Asp237, which is replaced by a neutral residue Asn236 in b^0,+^AT, indicating that one acidic residue is sufficient for recognizing cationic substrates in system b^0,+^.

Because b^0,+^AT also transports neutral amino acids, we next compared b^0,+^AT with LAT1 (SLC7A5), a large neutral amino acid transporter of the SLC7 family. The key acidic residue Asp233 of b^0,+^AT corresponds to Gly255 of LAT1 (Fig. 6f), which was shown to be important for recognizing large amino acids (*9*). With this small Gly255 residue, LAT1 forms a so-called distal pocket, which might accommodate large hydrophobic substrates. In b^0,+^AT, this distal pocket is disrupted (Fig. 6f) by the replacement of Val408 with a larger Tyr408, along with Asp233. Therefore, b^0,+^AT would not prefer bulky amino acids, unlike LAT1, which can recognize much bulkier analogs such as T_3_ and JPH203 (*40*).

Multiple sequence alignment of HATs, CATs (*39*) and GkApcT showed that the unwound regions of TM1 (residues 44–48 in b^0,+^AT) are highly conserved (Fig. S12a), whereas the equivalent region of TM6 (residues 233–237 in b^0,+^AT) varies substantially (Fig. S12b)(*39*). Asp233 and Asn236 in b^0,+^AT have drawn our attention, since at least one of these residues is negatively charged in systems b^0,+^, y^+^L and CATs, which all can transport cationic amino acids (Fig. S12b). To test the importance of these residues for substrate specificity, we mutated Asp233 and Asn236 and measured transport activities for cationic (Orn), neutral (Tyr) and small (Ala) amino acids. When Asp233 is mutated to neutral amino acids (D233G, D233I, and D233S), b^0,+^AT did not transport Orn (Fig. 6g), confirming that the positive charge is compensated by the negative charge of Asp233. By contrast, recognition of neutral amino acids did not fully require the negative charge of Asp233, as D233S retained Tyr transport activity (Fig. 6h). The N236D mutation mimics an Asp residue in y^+^L and CATs (Fig. S12b), and it retained Orn recognition to some degree but completely lost Tyr transport (Fig. 6g,h), supporting its importance for positive charge recognition in y^+^L and CATs. Kinetics of Orn and Tyr transport by these mutants (N236D and D233S) indicated slightly lower K_m_ as compared to WT, but largely lower V_max_ values, suggesting similar affinity but altered transport efficiency (Fig. 6j,k). The mutational effects were less prominent in the transport of small amino acids like Ala (Fig. 6i). These results demonstrate the importance of both Asp233 and Asn236 for the selectivity of cationic and large neutral amino acids, and that their functional properties are partially interchangeable among the SLC7 members. Moreover, the interchangeable properties of D233S and N236D imply that system b^0,+^ is more similar to system y^+^L than to system L or CATs, suggesting an evolutionary significance.

The system y^+^L transporters operate in a Na^+^-dependent manner when transporting neutral amino acids (*41*), whereas system b^0,+^ does not show any Na^+^ dependency. To investigate this behavior, we performed transport assays in the presence of Na^+^. Tyr transport by b^0,+^AT WT or D233S did not require Na^+^ (Fig. 6l), consistent with an Na^+^-independent system b^0,+^ (*39*). Although our single mutants D233S and N236D were designed to mimic system y^+^L and indeed showed functional properties to partly switch the substrate selectivity (Fig 6g,h), the double mutants (D233S/N236D and D233V/N236D) fully abolished the transport for all substrates (Fig. 6l). These results suggest an additional requirement of other residues for the full recognition of the substrate and Na^+^ in system y^+^L, such as residues in TM3 and TM8, which are also involved in substrate recognition. Thus, the molecular basis of Na^+^ dependence in the system of y^+^L transporters remains an open question.

## Discussion

### System b^0,+^ biogenesis and cystinuria

Although the super-dimerization of system b^0,+^ was reported many years ago (*42*), its biological relevance has so far remained unclear. Here, using structure-based biochemical assays, we have shown that super-dimerization is a key step for system b^0,+^ maturation (Fig. 7a–d). As N-glycan maturation is a Golgi-localized process, we propose that this super-dimerization precedes ER-Golgi trafficking, acting as a checkpoint for protein quality control (Fig. 7a). We also showed that super-dimerization is Ca^2+^ dependent, which is plausible given the rich Ca^2+^ pool of the ER lumen, where rBAT folding occurs (*11*) (Fig. 7a). Such a role of Ca^2+^ is reminiscent of its known regulatory roles in protein folding machineries and post-translational modifications (*32*), but the current case is unique in that Ca^2+^ binds directly to the protein being folded. Other parts of the structure also play important roles in system b^0,+^ biogenesis. For example, cystinuria mutants R365W and M467T have been known to facilitate proteolytic degradation (*11*) (Fig. 7b), and our structure indicates that these substitutions affect the domain A–TMD and A–C interfaces and probably destabilize the assembled system b^0,+^. Human rBAT possesses an additional N-glycan on Asn575’, which stabilizes the domain A–C interface and is indispensable in the maturation of other glycans (*23*). The C-terminal peptide of rBAT forms a β-hairpin that mediates interactions with b^0,+^AT and the lipid bilayer, which explains how the L678P mutant affects protein stability (*23*) (Fig. 7b). Even upon correct folding and cell-surface trafficking (Fig. 7c), some of the non-type I mutations such as V170M impair transport (Fig. 7d). Interestingly, our mutational analysis has shown that the b^0,+^AT G105R fails to fold, indicating that some of non-type I mutations are, at the molecular level, protein-folding defects (Fig. 7a).

**Figure 7.**
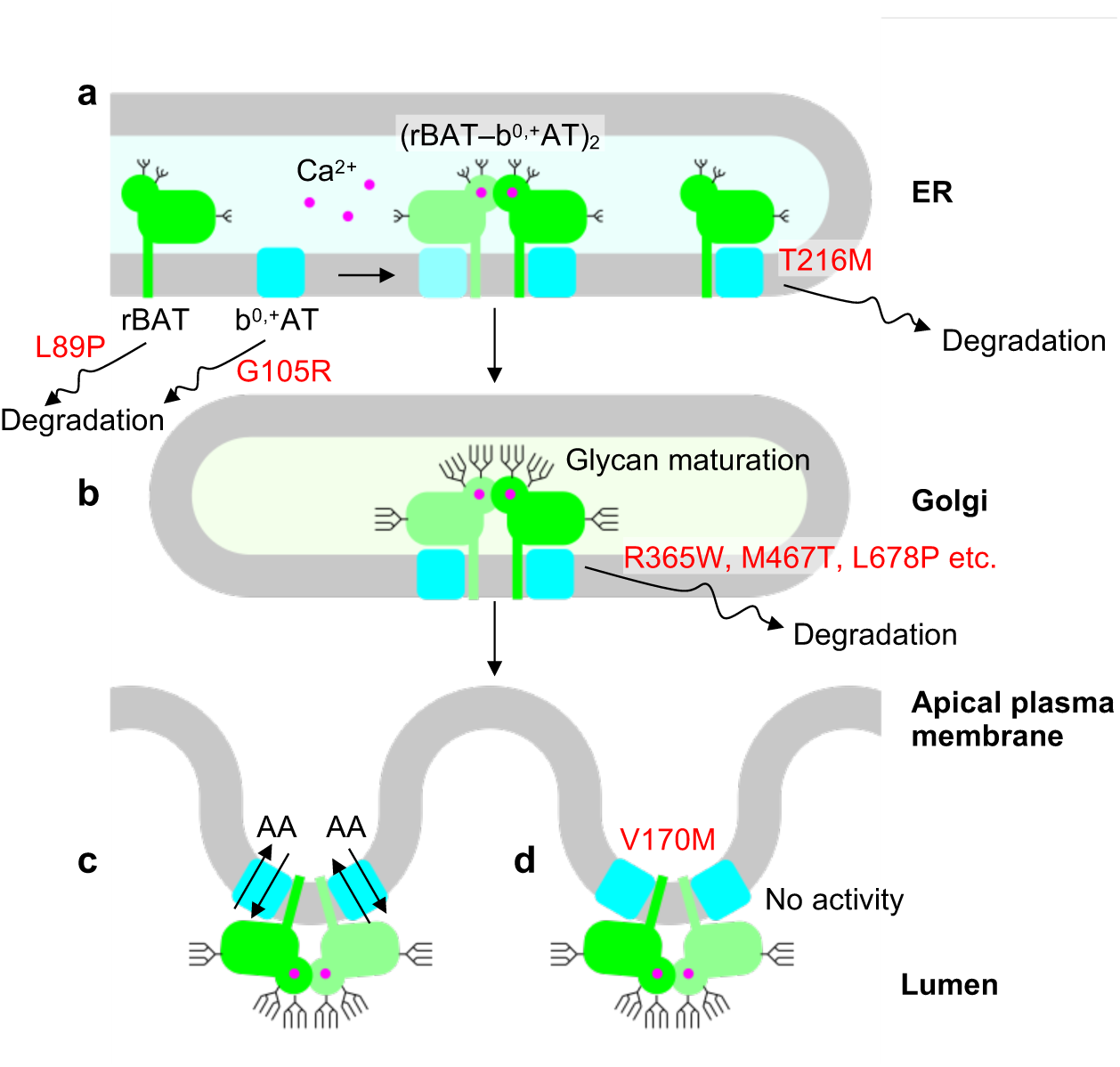
A working model for system b^0,+^ biogenesis and its defects in cystinuria. **a)** In the ER, rBAT and b^0,+^AT are folded, assemble into b^0,+^AT–rBAT and then super-dimerize upon binding of Ca^2+^. If super-dimerization is impaired, as in T216M or Ca^2+^-site mutants, the complex cannot exit the ER and will thus be degraded. Notably, since oxidative folding of rBAT requires association with b^0,+^AT [ref], and b^0,+^AT folding mutants (e.g., G105R) or rBAT assembly mutants (e.g., L89P) abolish initial assembly and thus lead to early degradation. **b)** In the Golgi, the super-dimeric b^0,+^AT–rBAT undergoes glycan maturation. Cystinuria mutants such as R365W and M467T can get partially maturated but are prone to degradation due to their lower stability. **c)** Upon correct maturation, super-dimeric b^0,+^AT–rBAT will be trafficked to the apical membrane of epithelial cells. Microvilli consist of numerous membrane protrusions, consistent with the curved lipid bilayer of b^0,+^AT–rBAT. **d)** Non-type I mutations (e.g., V170M) abolish transport activity.

LeuT-fold transporters are known to operate via a so-called ‘rocking-bundle’ mechanism, in which the bundle domain undergoes a rocking motion to mediate substrate transport (*43*). This major conformational change is associated with minor conformational changes of other structural elements, including EL4a, EL4b and IL1, which regulate the gating process. In our structure of b^0,+^AT, EL4a and EL4b form a lid above the extracellular gate, interacting with TM1, EL2 and the Aα4-β5 loop (Fig. S13a). A similar positioning of EL4a and EL4b is also seen in LAT1 (*9*), albeit with different interactions with CD98hc (Fig. S13b). The comparison of outward-facing and inward-facing structures of AdiC shows that, upon the inward-to-outward structural transition, EL4b dissociates from TM3, and EL4a undergoes upward movement to widen the transport pathway (Fig. S13c,d). We speculate that similar movements of EL4a and EL4b would be required for b^0,+^AT to transition to the outward-facing state. Such a movement explains how R365W affects the transport characteristics of b^0,+^AT (*44*) by altering the Aα4-β5 loop (Fig. 3a and Fig. S13a). A similar structural cross talk could also explain the incomplete negative dominance pattern of some of the non-type I cystinuria (*13*), where an inactive subunit inhibits the other subunit through the network of homo- and heteromeric interactions.

## Conclusions

Our study showed that b^0,+^AT–rBAT exists as a dimer of two heterodimers within the membrane and that this assembly is the key to system b^0,+^ maturation. Although membrane protein oligomerization has been widely observed in the contexts of protein stabilization (*45*), substrate translocation (*43*) or lipid ultrastructure (*46*), its obligatory requirement for N-glycan maturation is so far unique. The strict requirement might reflect the controlled trafficking of system b^0,+^ to the apical side of kidney brush border cells (*19*). Our study also revealed a previously unknown role of Ca^2+^ in super-dimerization and N-glycan maturation of system b^0,+^. As other possible factors that might control this process are currently unknown, further studies are needed to clarify possible links between the ER quality control, the Ca^2+^ dynamics and cystinuria. Finally, our study revealed that T216M, one of the most common cystinuria mutations worldwide (*1*), critically affects the higher-order assembly of the transporter and thereby causes type I cystinuria. Therefore, the restoration of super-dimerization may emerge as a potential therapeutic strategy for curing such types of cystinuria.

## Methods

### Protein expression and purification

Murine and ovine b^0,+^AT and rBAT genes were amplified from tissue cDNA (ZYAGEN) and cloned into the pEZT-BM vector (a gift from Ryan Hibbs; Addgene plasmid # 74099) (*47*). The cloned ovine rBAT sequence contained multiple nucleotide polymorphisms as compared to the closest UniProt entry (UniProt ID: W5P8K2). All substitutions are silent except for Arg181’ > Gln181’, which is a non-conserved residue on the rBAT surface and is not relevant for the structures presented in this study. For purification, rBAT was fused with the N-terminal His_8_-EGFP tag and a TEV cleavage site. b^0,+^AT was fused with the C-terminal TwinStrep II tag. Protein expression was performed as described (*47*). Briefly, P2 or P3 baculovirus was produced in Sf9 cells and used to transfect HEK293S GnTI^-^ suspension cells cultured in Freestyle 293 (Gibco) with 2% FBS at 37°C with 8% CO_2_. At the cell density of 2–3 × 10^6^ cells ml^-1^, the two concentrated viruses were added simultaneously with 5 mM sodium butyrate. After 12–18 hours, the culture temperature was decreased to 30°C, and cells were harvested 48 hours post-infection.

All purification procedures were performed at 4°C unless otherwise stated. Harvested cells were resuspended in lysis buffer (50 mM Tris, pH 8.0, 150 mM NaCl and cOmplete protein inhibitor cocktail) and lysed by sonication with a probe sonicator or by homogenization in a glass potter. The membrane fraction was pelleted by ultracentrifugation at 40,000 rpm, 1 h 10 min (45Ti, Beckman), resuspended in lysis buffer and stored in aliquots at –80°C until use. The membrane fraction was solubilized in lysis buffer supplemented with 0.5% lauryl maltose neopentyl glycol (LMNG) and 0.1% cholesterol hemisuccinate (CHS) for 3 hours. Insoluble material was removed by ultracentrifugation at 40,000 rpm, 30 min (45Ti, Beckman). The supernatant was incubated with GFP nanotrap resin (*48*) for 2 hours, and the resin was washed with ten column volumes of purification buffer (20 mM Tris-HCl, pH 8.0, 150 mM NaCl, 0.01% LMNG and 0.002% CHS). The target protein was cleaved off the column with TEV protease and subjected to size exclusion chromatography on a Superose 6 Increase 3.2/300 column connected to the Ettan system (GE Healthcare) operated at room temperature. The peak fraction was concentrated and used for further analyses (Fig. S2a).

For initial negative-stain EM and cryo-EM trials in LMNG, samples were prepared by slightly different protocols. For negative-stain EM, murine and ovine b^0,+^AT constructs were fused with Strep II instead of the TwinStrep II tag and the target complex was purified in two steps, first with the Strep XT resin and then with the GFP nanotrap. For initial cryo-EM in LMNG, b^0,+^AT fused with TwinStrep II was purified in a single-step with Strep XT resin and then subjected to size exclusion chromatography without cutting GFP.

### Liposome assay

Proteoliposomes were prepared by a published procedure (*9*). Briefly, soy PC (40% lecithin) was dissolved in chloroform and dried into a thin film under a continuous flow of nitrogen gas. The film was resuspended into liposome buffer (20 mM HEPES, pH 7.0 and 120 mM NaCl) at a lipid concentration of 20 mg ml^-1^ and sonicated in a bath sonicator for 30 min at room temperature to yield unilamellar vesicles. Purified proteins were added at a protein-to-lipid ratio of 1:166 (w/w) and reconstituted by three freeze-thaw cycles. Reconstituted proteoliposomes were diluted threefold and centrifuged at 10,000 rpm, 10 min to remove large aggregates, and then the supernatant was ultracentrifuged at 40,000 rpm, 2 h to pellet the liposomes. The pellet was stored at –80°C until subsequent experiments. For uptake experiments, the proteoliposome pellet was resuspended in reaction buffer (20 mM HEPES, pH 7.0 and 120 mM NaCl) with or without 1 mM □-Arg and sonicated with a probe sonicator to form unilamellar vesicles and load the counter-substrates into the liposomes. At this point, small samples were subjected to SDS-PAGE to check the reconstitution efficiency (Fig. S2c). Proteoliposomes were passed through a G-50 gel filtration column pre-equilibrated with reaction buffer to remove free □-Arg and then used immediately for transport assays.

To start the transport reactions, proteoliposomes were mixed with four volumes of extraliposomal solution (reaction buffer with 0.11 μM □-[^3^H]Arg; 54.5 Ci/mmol; PerkinElmer) and incubated at 25°C. At indicated time points, aliquots were taken and applied to Dowex 50WX4 cation exchange resin in a spin column to remove free □-[^3^H]Arg. The resin had been pre-equilibrated with reaction buffer and pre-chilled on ice to stop the reaction. After centrifugation at 700 ×g, 1 min, the eluted proteoliposomes were mixed with liquid scintillation cocktail (Rotiszint Eco Plus, Carl Roth) and radioactivity was measured with a liquid scintillation counter (Tri-carb 1500, Packard).

### Negative-stain electron microscopy

After the GFP nanotrap step, purified b^0,+^AT–rBAT samples were diluted to about 0.02 mg ml^-1^ and applied onto glow-discharged copper grids coated with continuous carbon film. Staining was performed with three drops of 1% uranyl formate and blotting with Whatman Grade 50 filter paper. LAT1–CD98hc was prepared as described (*9*) and stained similarly. Grids were imaged in a Tecnai BioTwin electron microscope operated at 120 kV. Roughly 50 images were recorded for each sample at a 49,000 × magnification with a 2.2 Å nominal pixel size and analyzed in RELION3.0 (*49*).

### Nanodisc reconstitution

Nanodisc scaffold protein MSPE3D1 was prepared as described (*50*). Briefly, MSPE3D1 with N-terminal His_7_ tag and a TEV cleavage site was expressed in *E. coli* BL21 cells. MSPE3D1 was purified on NiNTA column with four washes with buffer containing Triton X-100 and cholate, and eluted with 400 mM imidazole. The His tag was cleaved by TEV protease, and the sample was passed through a NiNTA column and then dialyzed against storage buffer (20 mM Tris, pH8.0 and 150 mM NaCl). Purified MSPE3D1 was concentrated and stored in aliquots at –80°C.

Nanodisc reconstitution was performed as described (*50*). Briefly, the lipid mixture was prepared by dissolving phosphatidylcholine (PC), phosphatidylglycerol (PG) and cholesterol (CHOL) in chloroform at a ratio of 2:2:1 (w/w). The mixture was dried into a thin film in a glass vial under nitrogen gas flow and then resuspended in 50 mM sodium cholate solution to a final phospholipid concentration of 20 mg ml^-1^. Purified b^0,+^AT–rBAT was mixed with MSP1E3D1 and the lipid mixture to a protein:scaffold:lipid ratio of 1:5:200 (mol/mol) adding 20 mM sodium cholate, and incubated at 4°C with gentle end-to-end rotation. After incubation for 1 h, a first batch of Bio-Beads SM-2 resin (500 mg per 1 ml mixture) was added to start detergent removal. After 2 h, a second batch was added, and the detergent removal proceeded overnight. The reconstituted nanodiscs were recovered by passing through a spin column and subjected to size exclusion chromatography on a Superose 6 Increase 3.2/300 column at room temperature. Peak fractions were concentrated to about 10 mg ml^-1^ and immediately used for cryo-EM grid preparation.

### Sample vitrification for cryo-EM

All cryo-EM grids used in this study were Quantifoil 400 copper R1.2/1.3 grids. Grids were pre-washed in acetone and dried for at least 1 h and then glow-discharged twice in a Pleco glow discharger under 0.4 mBar air at 15 mA for 45 seconds. For vitrification, a Vitrobot Mark I was set to 6°C and 100% humidity, and filter paper for blotting (Whatman Grade 595) was pre-equilibrated in the chamber for about 1 h. 3 μl of protein solution at 10 mg ml^-1^ were applied to the grids, blotted for 3–4 seconds with blot force 20, and plunge-frozen. To improve sample distribution, 1.5 mM fluorinated Fos-Choline-8 was added to the sample immediately before vitrification. For detergent and nanodisc dataset #1, we added 10 mM substrate Arg about 30 min before vitrification, hoping to visualize the bound substrate, but none of the resulting maps showed any substrate density, indicating that the protein was in the apo state.

### Cryo-EM data collection

All cryo-EM data were recorded on Titan Krios microscopes operated at 300 kV. For b^0+^AT–rBAT in LMNG, the data were recorded on Titan Krios G3i at 105,000 × magnification with a K3 direct electron detector operated in a counting mode, resulting in a calibrated pixel size of 0.837 Å. The electron exposure rate was set to 15 e pix^-1^ sec^-1^, and the movies were recorded for 2 seconds with 50 frames, resulting in a total exposure of 50 e Å^-2^ on the specimen. For b^0,+^AT–rBAT in nanodisc, data were recorded in three separate sessions. In first session, settings were as for the LMNG dataset. In the second session, similar settings were used on another Titan Krios G2 microscope, equipped with a K3 camera with a calibrated pixel size of 0.831 Å. In third session, a Titan Krios G2 with Falcon III camera was used. For the Falcon III data collection, the magnification was set to 96,000 ×, resulting in a calibrated pixel size of 0.837 Å, and the exposure rate was set to 1.5 e pix^-1^ sec^-1^. Each movie was recorded for 30 seconds with 50 frames in a counting mode, resulting in a total exposure of 40 e Å^-2^. All movies were saved in gain-normalized TIFF format as specified by the EPU software (Thermo Fischer Scientific).

### Image processing

Single-particle image processing was performed in RELION 3.0 (*51*) (Fig. S3a,b). For the nanodisc datasets, data from three sessions were processed separately to the particle polishing step. Beam-induced motion was estimated and corrected in MotionCor2 (*52*) using 5 × 5 patches with dose-weighting. Defocus and astigmatism were estimated in Gctf4 (*53*). Initial particle sets were picked by using a Laplacian-of-Gaussian filter from a few dozen micrographs and extracted with a box size of 330 pix down-sampled to 80 pix. After 2D and 3D classifications, the resulting 3D map served as a template for automated picking in RELION. In parallel, coordinates from the resulting 3D classes were used to train Topaz (*54*) for neural network-based picking. The Topaz network was trained on dataset #1 with 8 × down-sampled micrographs and around 6,000 input coordinates. Particles picked by the two programs were extracted separately with down-sampling from 330 pix to 80 pix and subjected to rounds of independent 3D classification. After removing duplicate particles, good particles were re-extracted with a box size of 420 pix down-sampled to 320 pix, resulting in 1.099 Å/pix for nanodisc dataset #1. Particles were refined with an auto-refine procedure with a soft mask and C2 symmetry imposed. After the initial 3D refinement, particle trajectories and weighting B/k-factors were refined by Bayesian polishing (*55*), and then per-particle defocus values were refined as implemented in RELION 3.0 (*51*). For the detergent dataset, CTF-refined particles yielded the final reconstruction at 3.91 Å resolution according to the FSC = 0.143 criterion (Fig. S3a,d).

For the nanodiscs datasets, CTF-refined particles were further processed in RELION 3.1, to accommodate the different pixel sizes in different datasets (*56*). Particles from the three datasets were merged with different rlnOpticsGroup labels, and then refined against the reference map of dataset #1 (Fig. S3c). Resulting alignments and volumes were used to correct possible errors in pixel sizes by using anisotropic magnification correction, which reported less than 0.5% errors in the assumed pixel sizes. The 3D auto-refinement from the merged particles after magnification correction with C2 symmetry yielded a 2.88 Å map of the full complex in lipid nanodisc (Fig. S3c–e).

The map of the full complex indicated structural flexibility between the two b^0,+^AT– rBAT protomers. To improve local map quality, we took two approaches: (i) focused refinement for the rBAT ectodomain and (ii) multi-body refinement of individual b^0,+^AT–rBAT subcomplexes. For focused refinement, the TMD signal was subtracted from individual particles, and then subtracted particles were refined against the rBAT ectodomain map with C2 symmetry imposed, yielding a 2.68 Å resolution map for the rBAT ectodomain (Fig. S3e). Multi-body refinement was performed with two bodies, each covering an individual b^0,+^AT–rBAT protomer and half the micelle. Bayesian priors were set to 10 degrees for body orientations and 3 pixels for translations. After multi-body refinement, we merged the two bodies to obtain a single reconstruction for the b^0,+^AT–rBAT heterodimer subcomplex, since both bodies consist of identical sets of subunits. To do this, signal subtraction was performed for each body as described (*57*), and the resulting particles were merged and subjected to rounds of 3D classification to further select good particles, and refined against a map of the b^0,+^AT– rBAT heterodimer. Final refinement yielded a 3.05 Å resolution map for the b^0,+^AT– rBAT heterodimer subcomplex with improved local resolution for the TMD (Fig. S3f).

To visualize the major conformational heterogeneity in the dataset, we performed a principal component (PC1) analysis. We then divided the particles into three subsets of roughly equal size along the PC1 axis and performed normal 3D refinement for each subset. Morphing of the resulting three maps revealed a flexible motion of two b^0,+^AT molecules relative to the more rigid rBAT–rBAT assembly (Movie S1).

### Model building and refinement

The initial model for b^0,+^AT–rBAT was generated by homology modelling using MODELLER 9.22 (*58*). Homologous proteins were first identified using BLAST, and top hits were selected to cover the entire sequence of rBAT. Three structure were thus selected and used for homology modelling, namely *Bacillus cereus* oligo-1,6-glucosidase (PDB 1UOK), *Xanthomonas campestris* alpha-glucosyl transfer enzyme (PDB 6AAV) and LAT1–CD98hc (PDB 6IRS). The input multiple sequence alignment generated in Clustal Omega (*59*) covered almost all regions except for the C-terminal peptide of rBAT (residues 655–685). The MODELLER parameters were set as follows: library_schedule = autosched.slow; max_var_iterations = 300; md_level = refine.slow; repeat_optimization = 2; starting_model = 1; and ending_model = 5. All five resulting models MODELLER showed good scores with slightly different orientations between ED and TMD. One of them was used for further model building.

All manual model building was performed in COOT 0.8.9 (*60*). The cryo-EM maps resolved all residues except for the disordered terminal residues (residues 1–28 and 484–487 for b^0,+^AT and 1–62 for rBAT). After building the protein, prominent densities on the predicted N-glycosylation sites were built as acetyl glucosamine units (GlcNAc). A spherical density within the domain B was built as a Ca^2+^ ion (see Main Text). Among the numerous lipid densities resolved around the TMD, two flat-shaped densities in the outer leaflet were assigned as cholesterols. One density near TM1’, which had two elongated tails, was modelled as a phosphatidylcholine. Both lipids had been included in the lipid nanodisc composition (see Nanodisc reconstitution in Methods). Ligand restraints were generated in Acedrg (*61*) and applied throughout manual model building and refinement.

All model refinements were performed in phenix.real_space_refine in PHENIX (version 1.16-dev3689) (*62*). After iterative refinement of b^0,+^AT–rBAT against the heterodimer map, the model of the rBAT ectodomain was independently refined against the rBAT ectodomain map with a refinement resolution set to 2.6 Å. The refined rBAT ectodomain model was then merged back to the heterodimer model and refined again. For the full complex map, the two heterodimer models were fitted as rigid bodies. Model validation was performed with MolProbity (*63*) Validation statistics are shown in Table 1.

For structure visualization, UCSF ChimeraX and the associated software packages were used (*64*). The electrostatic potentials were calculated with APBS (*65*). Multiple sequence alignments were generated with Clustal Omega (*59*), and formatted with ESPript 3 (*66*).

### Endo H assay

The Endo H assay was performed as described previously (*11*) with small modifications. One modification was that we used fluorescent tags instead of using Western blotting (*11*). This not only simplified the experimental procedures, but also allowed dual-color fluorescence detection to visualize the heteromeric complexes on SDS-PAGE gels directly. For the assay, the pEZT-BM expression plasmids as described above were adopted with minimal modification. The rBAT plasmid was identical to that used for structural analysis (with the N-terminal GFP tag; pEZT-NGFP-OaSLC3A1 plasmid) and the b^0,+^AT was fused with an additional mCherry tag on the C-terminus after TwinStrep II (the pEZT-C-TwinStrep-mCherry-OaSLC7A9 plasmid).

Protein co-expression was performed in HeLa cells cultured at 37°C, 8% CO_2_ in DMEM supplemented with 10% FBS (Gibco) and penicillin/streptomycin mixture (Pen Strep, Gibco). Two days before transfection, cells were seeded onto 24-well plates at 0.3 × 10^6^ cells/well. The culture medium was exchanged with 500 μl of fresh DMEM with 10% FBS and Pen Strep approximately 10 min before transfection. For transient co-transfection, 250 ng each of two plasmids coding b^0,+^AT and rBAT were mixed into 100 μ of DMEM, and 1.5 μl of Polyjet reagent in 100 μl DMEM were added. After incubation at room temperature for 15–20 min, DNA:polyjet mixture was added dropwise to the HeLa cell culture, and expression proceeded at 37°C, 8% CO_2_. After 12–18 h post transfection, the medium was exchanged with 1 ml of DMEM with 10% FBS and Pen Strep. Cells were harvested at 48 h post transfection, pelleted at 5,000 rpm for 1 min, and stored at –20°C for subsequent analysis.

Pelleted cells were thawed and washed once with 500 μl lysis buffer (50 mM Tris-HCl, pH 8.0, 150 mM NaCl and cOmplete protease inhibitor), re-pelleted, and re-suspended in 100 μl lysis buffer. Cells were lysed by brief sonication with a probe sonicator (0.5 sec pulse and 2 seconds interval, 10 cycles) and the cell lysate was directly used for the assay. The glycosidase reaction was performed with Endo H enzyme (NEB) under non-denaturing conditions following manufacturer’s instructions. Briefly, 5 μl of cell lysate was added with 0.2 μl Endo H enzyme and topped to 7 μl with MilliQ water and 10 × Glyco 3 buffer. The reaction was performed at 37°C for 5 h. For negative controls, Endo H enzyme was replaced with MilliQ water. After the reaction, the reactant was mixed with the 2 × SDS-PAGE sample buffer containing 100 mM β-mercaptoethanol and run on 12% Mini-PROTEAN TGX Precast Gel (Bio-Rad) without a boiling step. As a molecular-weight marker, 1.5 μl of pre-stained Page Ruler (Thermo Scientific) was run in parallel. The gels were directly imaged on Gel Analyzer (Bio-Rad) without a fixation step. Fluorescent wavelengths and exposure times were set as follows: EGFP (488 nm, 120 sec), mCherry (546 nm, 1440 sec) and pre-stained marker (680 nm, 2 sec).

### Site-specific Cys cross-linking assay

Protein co-expression and cell lysis were performed as in the Endo H assay. To prevent spontaneous reduction of disulfide bonds after cell lysis, lysed cells were first treated with 0.5 mM 4-DPS at 4 °C for 2 min to generate oxidizing conditions. Then, to prevent non-specific disulfide bond formation after SDS denaturation, free cysteines were blocked by treating with 500 mM iodoacetamide at 4 °C for 5 min. The treated lysates were mixed with the 2 × SDS-PAGE sample buffer with or without 100 mM β-mercaptoethanol and run on 12% Mini-PROTEAN TGX Precast Gel (Bio-Rad) without a boiling step. Gels were imaged in the same way as in the Endo H assay.

### Localization of rBAT and b^0,+^AT

Localization of rBAT and b^0,+^AT was examined in HeLa cells. Cells (1.6 x 10^5^ cells/well) were seeded on collagen-coated coverslips in a 6-well plate and cultured in MEM containing 10% FBS at 37 □ and 5% CO_2_. Transfection was performed at 24 hours after seeding. Wildtype or mutant clones of rBAT (the pEZT-NGFP-OaSLC3A1 plasmid) and wildtype b^0,+^AT (the pEZT-C-TwinStrep-mCherry-OaSLC7A9 plasmid) were transfected into HeLa cells at 1:1 molar ratio by using PEI MAX (40,000 MW Linear; Polysciences, Inc.). The ratio of DNA : PEI was 4 µg total DNA : 8 µg PEI for each well. Cells were continuously cultured for 2 days. For imaging, the cells were washed with PBS and fixed with 4% w/v paraformaldehyde in PBS for 15 min at room temperature. Cells were then washed with PBS and incubated with DAPI (1 ng) in PBS for 30 min. After the PBS wash, specimens were mounted with Fluoromount (Diagnostic BioSystems), and imaged with a Keyence BZ-X710 microscope. Data were processed using ImageJ ver. 1.51 (NIH).

### Transport assay in transfected cell lines

For cell-based transport experiments, rBAT and b^0,+^AT genes were cloned into the pEG BacMam vector (a gift from Eric Gouaux (*67*)) without a tag. Transport functions of rBAT and b^0,+^AT mutants were measured in HEK293 cells. The cells were seeded on poly-D-lysine-coated 24-well plate at 1.7 x 10^5^ cells/well and cultured in DMEM containing 10% FBS, at 37 □ and 5% CO_2_. Transfection was performed at 24 hours after seeding. Wildtype or mutant clones of rBAT (the pEG-OaSLC3A1 plasmid) and b^0,+^AT (the pEG-OaSLC7A9 plasmid) were transfected at 1:1 molar ratio into HEK293 cells by using lipofectamine 3000 (Thermo) following the manufacturer’s protocol. After transfection, cells were cultured for 2 days. Transport assays were performed as described (*68*). Briefly, transport of 100 μM radioisotope substrates (L-[^14^C]ornithine (2 Ci/mol; Moravek Biochemicals), L-[^3^H]tyrosine (10 Ci/mol; PerkinElmer) and L-[^3^H]alanine (5 Ci/mol; Moravek Biochemicals)) were measured for 3 min in Na^+^-free HBSS pH 7.4. Transport of L-[^3^H]tyrosine was assayed in the presence of 10 μM SKN103 to inhibit endogenous LAT1 function (*69*). For uptake of L-[^3^H]tyrosine in Na^+^-containing condition, HBSS buffer was used. After terminating the reaction and cell lysis, an aliquot of the lysate was used to measure protein concentration by BCA protein assay (Takara Bio). The lysate was mixed with Optiphase Hisafe 3 (PerkinElmer), and radioisotope activity was monitored by LSC-8000 β-scintillation (Hitachi). Data shown in the figures were those subtracted by the uptake values in Mock cells (the cells transfected with empty plasmids). Kinetics of L-[^14^C]ornithine and L-[^3^H]tyrosine transport were determined for 3 min using radioisotope substrates at a concentration of 20 – 400 μM. Curves were fitted to Michaelis–Menten plots (GraphPad Prism 8.4).

### Data availability

The atomic coordinates have been deposited to Protein Data Bank under accession numbers 7NF6 (b^0,+^AT–rBAT heterodimer), 7NF7 (rBAT ectodomain) and 7NF8 (full complex composite model). Cryo-EM maps have been deposited to Electron Microscopy Data Bank under accession numbers EMD-12296 (b^0,+^AT–rBAT heterodimer), EMD-12297 (rBAT ectodomain) and EMD-12298 (full complex). All other data will be available upon request.

## Acknowledgments

We thank Susann Kaltwasser, Simone Prinz, Mark Linder and Sonja Welsch in the Central Electron Microscopy Facility of Max Planck Institute of Biophysics for technical assistance in electron microscopy; Sabine Häder, Christina Kunz and Heidi Betz for help in lab experiments; Juan Castillo, Özkan Yildiz, the Central IT team and the Max Planck Computing and Data Facility for maintaining the computational infrastructure; David Wöhlert and Martin Centola for technical advice and help in mammalian cell culture and protein expression; Department of Molecular Membrane Biology, Max Planck Institute of Biophysics for sharing HeLa cell culture; Yoko Tanaka for technical assistance in transport assays; and Pornparn Kongpracha for preliminary proteomics analysis. This work was supported by the Max Planck Society to Y.L, D.J.M. and W.K., AMED under Grant Numbers JP21gm0810010 and JP21lk0201112 and JSPS KAKENHI (JP21H03365) to S.N., and MEXT/JSPS KAKENHI (JP19K07373) and Mochida Memorial Foundation grant for Medical and Pharmaceutical research to P.W. Y.L was supported by Toyobo Biotechnology Foundation Fellowship and Human Frontier Science Program Long-Term Fellowship.

## Author contributions

Y.L. and S.N. initiated the study. Y.L. performed the structural studies and *in vitro* biochemical experiments. P.W. performed the cell-based transport assays. P.W. and S.M. performed fluorescence imaging. D.M. maintained and aligned the electron microscopes and directed data collection. S.N. and W.K. directed and supervised the project. Y.L. P.W. and S.N. wrote the manuscript, with contributions from W.K.

## References

1. J. Chillarón, M. Font-Llitjós, J. Fort, A. Zorzano, D. S. Goldfarb, V. Nunes, M. Palacín, Pathophysiology and treatment of cystinuria. Nat Rev Nephrol. 6, 424–434 (2010).

2. K. Thomas, K. Wong, J. Withington, M. Bultitude, A. Doherty, Cystinuria—a urologist’s perspective. Nat Rev Urol. 11, 270–277 (2014).

3. D. Joly, P. Rieu, A. Méjean, M.-F. Gagnadoux, M. Daudon, P. Jungers, Treatment of cystinuria. Pediatr Nephrol. 13, 945–950 (1999).

4. T. Zee, N. Bose, J. Zee, J. N. Beck, S. Yang, J. Parihar, M. Yang, S. Damodar, D. Hall, M. N. O’Leary, A. Ramanathan, R. R. Gerona, D. W. Killilea, T. Chi, J. Tischfield, A. Sahota, A. Kahn, M. L. Stoller, P. Kapahi, α-Lipoic acid treatment prevents cystine urolithiasis in a mouse model of cystinuria. Nat Med. 23, 288–290 (2017).

5. M. Calonge, P. Gasparini, J. Chillarón, M. Chillón, M. Gallucci, F. Rousaud, L. Zelante, X. Testar, B. Dallapiccola, F. Silverio, P. Barceló, X. Estivill, A. Zorzano, V. Nunes, M. Palacín, Cystinuria caused by mutations in rBAT, a gene involved in the transport of cystine. Nat Genet. 6, 420–425 (1994).

6. L. Feliubadaló, M. Font, J. Purroy, F. Rousaud, X. Estivill, V. Nunes, E. Golomb, M. Centola, I. Aksentijevich, Y. Kreiss, B. Goldman, M. Pras, D. L. Kastner, E. Pras, P. Gasparini, L. Bisceglia, E. Beccia, M. Gallucci, L. de Sanctis, A. Ponzone, G. F. Rizzoni, L. Zelante, M. T. Bassi, A. L. George, M. Manzoni, A. D. Grandi, M. Riboni, J. K. Endsley, A. Ballabio, G. Borsani, N. Reig, E. Fernández, R. Estévez, M. Pineda, D. Torrents, M. Camps, J. Lloberas, A. Zorzano, M. Palacín, I. C. Consortium, Non-type I cystinuria caused by mutations in SLC7A9, encoding a subunit (bo,+AT) of rBAT. Nat Genet. 23, 52–57 (1999).

7. J. Chillarón, R. Roca, A. Valencia, A. Zorzano, M. Palacín, Heteromeric amino acid transporters: biochemistry, genetics, and physiology. American journal of physiology. Renal Physiol. 281, F995–1018 (2001).

8. R. Yan, X. Zhao, J. Lei, Q. Zhou, Structure of the human LAT1–4F2hc heteromeric amino acid transporter complex. Nature. 568, 127–130 (2019).

9. Y. Lee, P. Wiriyasermkul, C. Jin, L. Quan, R. Ohgaki, S. Okuda, T. Kusakizako, T. Nishizawa, K. Oda, R. Ishitani, T. Yokoyama, T. Nakane, M. Shirouzu, H. Endou, S. Nagamori, Y. Kanai, O. Nureki, Cryo-EM structure of the human L-type amino acid transporter 1 in complex with glycoprotein CD98hc. Nat Struct Mol Biol. 26, 510–517 (2019).

10. M. Gabriško, Š. Janeček, Looking for the ancestry of the heavy chain subunits of heteromeric amino acid transporters rBAT and 4F2hc within the GH13 □ amylase family. FEBS Journal. 276, 7265–7278 (2009).

11. P. Bartoccioni, M. Rius, A. Zorzano, M. Palacín, J. Chillarón, Distinct classes of trafficking rBAT mutants cause the type I cystinuria phenotype. Hum Mol Genet. 17, 1845–1854 (2008).

12. Y. Shigeta, Y. Kanai, A. Chairoungdua, N. Ahmed, S. Sakamoto, H. Matsuo, D. K. Kim, M. Fujimura, N. Anzai, K. Mizoguchi, T. Ueda, K. Akakura, T. Ichikawa, H. Ito, H. Endou, A novel missense mutation of SLC7A9 frequent in Japanese cystinuria cases affecting the C-terminus of the transporter. Kidney Int. 69, 1198–1206 (2006).

13. M. Font-Llitjós, M. Jiménez-Vidal, L. Bisceglia, D. M. Perna, L. de Sanctis, F. Rousaud, L. Zelante, M. Palacín, V. Nunes, New insights into cystinuria: 40 new mutations, genotype–phenotype correlation, and digenic inheritance causing partial phenotype. J Med Genet. 42, 58–68 (2005).

14. R. Yan, Y. Li, Y. Shi, J. Zhou, J. Lei, J. Huang, Q. Zhou, Cryo-EM structure of the human heteromeric amino acid transporter b^0,+^AT-rBAT. Sci Adv. 6, eaay6379 (2020).

15. D. Wu, T. N. Grund, S. Welsch, D. J. Mills, M. Michel, S. Safarian, H. Michel, Structural basis for amino acid exchange by a human heteromeric amino acid transporter. Proc Natl Acad Sci 117, 21281–21287 (2020).

16. T. Nakane, D. Kimanius, E. Lindahl, S. H. Scheres, Characterisation of molecular motions in cryo-EM single-particle data by multi-body refinement in RELION. eLife. 7, e36861 (2018).

17. M. Palacín, Y. Kanai, The ancillary proteins of HATs: SLC3 family of amino acid transporters. Pflügers Archiv. 447, 490–494 (2004).

18. H. Krishnamurthy, C. L. Piscitelli, E. Gouaux, Unlocking the molecular secrets of sodium-coupled transporters. Nature. 459, 347–355 (2009).

19. M. Furriols, J. Chillarón, C. Mora, A. Castelló, J. Bertran, M. Camps, X. Testar, S. Vilaró, A. Zorzano, M. Palacín, rBAT, related to L-cysteine transport, is localized to the microvilli of proximal straight tubules, and its expression is regulated in kidney by development. J Biol Chem. 268, 27060–27068 (1993).

20. S. Sakamoto, A. Chairoungdua, S. Nagamori, P. Wiriyasermkul, K. Promchan, H. Tanaka, T. Kimura, T. Ueda, M. Fujimura, Y. Shigeta, Y. Naya, K. Akakura, H. Ito, H. Endou, T. Ichikawa, Y. Kanai, A novel role of the C-terminus of b0,+AT in the ER–Golgi trafficking of the rBAT–b0,+AT heterodimeric amino acid transporter. Biochem J. 417, 441–448 (2009).

21. M. Rius, J. Chillarón, Carrier Subunit of Plasma Membrane Transporter Is Required for Oxidative Folding of Its Helper Subunit. J Biol Chem. 287, 18190–18200 (2012).

22. G. J. Peter, A. Davies, P. W. Watt, J. Birrell, P. M. Taylor, Interactions between the thiol-group reagent N-ethylmaleimide and neutral and basic amino acid transporter-related amino acid transport. Biochem J. 343, 169–176 (1999).

23. M. Rius, L. Sala, J. Chillarón, The role of N-glycans and the C-terminal loop of the subunit rBAT in the biogenesis of the cystinuria-associated transporter. Biochem J. 473, 233–244 (2016).

24. K. P. Chai, N. F. B. Othman, A.-H. Teh, K. L. Ho, K.-G. Chan, M. S. Shamsir, K. M. Goh, C. L. Ng, Crystal structure of Anoxybacillus α-amylase provides insights into maltose binding of a new glycosyl hydrolase subclass. Sci Rep. 6, 23126 (2016).

25. E. Boel, L. Brady, A. M. Brzozowski, Z. Derewenda, G. G. Dodson, V. J. Jensen, S. B. Petersen, H. Swift, L. Thim, H. F. Woldike, Calcium binding in .alpha.-amylases: an x-ray diffraction study at 2.1-.ANG. resolution of two enzymes from Aspergillus. Biochemistry. 29, 6244–6249 (1990).

26. M. Machius, N. Declerck, R. Huber, G. Wiegand, Activation of Bacillus licheniformis α-amylase through a disorder→order transition of the substrate-binding site mediated by a calcium–sodium–calcium metal triad. Structure. 6, 281–292 (1998).

27. Z. Li, W. F. Stafford, M. Bouvier, The metal ion binding properties of calreticulin modulate its conformational flexibility and thermal stability. Biochemistry. 40, 11193–11201 (2001).

28. A. Brockmeier, D. B. Williams, Potent lectin-independent chaperone function of calnexin under conditions prevalent within the lumen of the endoplasmic reticulum. Biochemistry. 45, 12906–12916 (2006).

29. S. K. Nigam, A. L. Goldberg, S. Ho, M. F. Rohde, K. T. Bush, S. MYu, A set of endoplasmic reticulum proteins possessing properties of molecular chaperones includes Ca(2+)-binding proteins and members of the thioredoxin superfamily. J Biol Chem. 269, 1744–1749 (1994).

30. H. K. Lamb, C. Mee, W. Xu, L. Liu, S. Blond, A. Cooper, I. G. Charles, A. R. Hawkins, The affinity of a major Ca2+ binding site on GRP78 is differentially enhanced by ADP and ATP. J Biol Chem. 281, 8796–8805 (2006).

31. H. A. Lucero, B. Kaminer, The role of calcium on the activity of ERcalcistorin/protein-disulfide isomerase and the significance of the C-terminal and its calcium binding. A comparison with mammalian protein-disulfide isomerase. J Biol Chem. 274, 3243–3251 (1999).

32. H. Coe, M. Michalak, Calcium binding chaperones of the endoplasmic reticulum. Gen Physiol Biophys. 28, F96–F103 (2009).

33. W. Auiewiriyanukul, W. Saburi, K. Kato, M. Yao, H. Mori, Function and structure of GH13_31 α-glucosidase with high α□(1→4) □glucosidic linkage specificity and transglucosylation activity. FEBS Lett. 592, 2268–2281 (2018)

34. M. Palacín, C. Mora, J. Chillarón, M. J. Calonge, R. Estévez, D. Torrents, X. Testar, A. Zorzano, V. Nunes, J. Purroy, X. Estivill, P. Gasparini, L. Bisceglia, L. Zelante, The molecular basis of cystinuria: the role of the rBAT gene. Amino Acids. 11, 225–246 (1996).

35. M. A. Font, L. Feliubadaló, X. Estivill, V. Nunes, E. Golomb, Y. Kreiss, E. Pras, L. Bisceglia, A. P. d’Adamo, L. Zelante, P. Gasparini, M. T. Bassi, A. L. George, M. Manzoni, M. Riboni, A. Ballabio, G. Borsani, N. Reig, E. Fernández, A. Zorzano, J. Bertran, M. Palacín, I. C. Consortium, Functional analysis of mutations in SLC7A9, and genotype–phenotype correlation in non-Type I cystinuria. Hum Mol Genet. 10, 305–316 (2001).

36. L. R. Forrest, R. Krämer, C. Ziegler, The structural basis of secondary active transport mechanisms. Biochim Biophys Acta Bioenerg. 1807, 167–188 (2010).

37. L. Ponzoni, S. Zhang, M. Cheng, I. Bahar, Shared dynamics of LeuT superfamily members and allosteric differentiation by structural irregularities and multimerization. Phil Trans R Soc B. 373, 20170177 (2018).

38. K. E. J. E. Jungnickel, J. L. Parker, S. Newstead, Structural basis for amino acid transport by the CAT family of SLC7 transporters. Nat Commun. 9, 550 (2018).

39. F. Verrey, E. I. Closs, C. A. Wagner, M. Palacin, H. Endou, Y. Kanai, CATs and HATs: the SLC7 family of amino acid transporters. Pflügers Archiv. 447, 532–542 (2004).

40. H.-C. Chien, C. Colas, K. Finke, S. Springer, L. Stoner, A. A. Zur, B. Venteicher, J. Campbell, C. Hall, A. Flint, E. Augustyn, C. Hernandez, N. Heeren, L. Hansen, A. Anthony, J. Bauer, D. Fotiadis, A. Schlessinger, K. M. Giacomini, A. A. Thomas, Reevaluating the Substrate Specificity of the L-Type Amino Acid Transporter (LAT1). J Med Chem. 61, 7358–7373 (2018).

41. R. Devés, C. Boyd, Transporters for cationic amino acids in animal cells: discovery, structure, and function. Physiol Rev. 78, 487–545 (1998).

42. E. Fernández, M. Jiménez-Vidal, M. Calvo, A. Zorzano, F. Tebar, M. Palacín, J. Chillarón, The structural and functional units of heteromeric amino acid transporters. The heavy subunit rBAT dictates oligomerization of the heteromeric amino acid transporters. J Biol Chem. 281, 26552–26561 (2006).

43. D. Drew, O. Boudker, Shared molecular mechanisms of membrane transporters. Annu Rev Biochem. 85, 1–30 (2015).

44. Pineda, C. A. Wagner, A. Bröer, P. A. Stehberger, S. Kaltenbach, J. L. Gelpí, R. M. D. Río, A. Zorzano, M. Palacín, F. Lang, S. Bröer, Cystinuria-specific rBAT(R365W) mutation reveals two translocation pathways in the amino acid transporter rBAT-b0,+AT. Biochem J. 377, 665–674 (2004).

45. K. Gupta, J. A. Donlan, J. T. Hopper, P. Uzdavinys, M. Landreh, W. B. Struwe, D. Drew, A. J. Baldwin, P. J. Stansfeld, C. V. Robinson, The role of interfacial lipids in stabilizing membrane protein oligomers. Nature. 541, 421–424 (2017).

46. T. B. Blum, A. Hahn, T. Meier, K. M. Davies, W. Kühlbrandt, Dimers of mitochondrial ATP synthase induce membrane curvature and self-assemble into rows. Proc Natl Acad Sci. 116, 4250–4255 (2019).

47. C. L. Morales-Perez, C. M. Noviello, R. E. Hibbs, Manipulation of Subunit Stoichiometry in Heteromeric Membrane Proteins. Structure. 24, 797–805 (2016).

48. U. Rothbauer, K. Zolghadr, S. Muyldermans, A. Schepers, C. M. Cardoso, H. Leonhardt, A versatile nanotrap for biochemical and functional studies with fluorescent fusion proteins. Mol Cell Proteomics. 7, 282–289 (2008).

49. .J. Zivanov, T. Nakane, B. O. Forsberg, D. Kimanius, W. J. Hagen, E. Lindahl, S. H. Scheres, New tools for automated high-resolution cryo-EM structure determination in RELION-3. eLife. 7, e42166 (2018).

50. F. Hagn, M. L. Nasr, G. Wagner, Assembly of phospholipid nanodiscs of controlled size for structural studies of membrane proteins by NMR. Nat Protoc. 13, 79–98 (2018).

51. J. Zivanov, T. Nakane, B. O. Forsberg, D. Kimanius, W. J. Hagen, E. Lindahl, S. H. Scheres, New tools for automated high-resolution cryo-EM structure determination in RELION-3. eLife. 7, e42166 (2018).

52. S. Q. Zheng, E. Palovcak, J.-P. P. Armache, K. A. Verba, Y. Cheng, D. A. Agard, MotionCor2: anisotropic correction of beam-induced motion for improved cryo-electron microscopy. Nat Methods. 14, 331–332 (2017).

53. K. Zhang, Gctf: Real-time CTF determination and correction. J Struct Biol. 193, 1–12 (2016).

54. T. Bepler, A. Morin, M. Rapp, J. Brasch, L. Shapiro, A. J. Noble, B. Berger, Positive-unlabeled convolutional neural networks for particle picking in cryo-electron micrographs. Nat Methods. 16, 1–8 (2019).

55. J. Zivanov, T. Nakane, S. H. W. Scheres, A Bayesian approach to beam-induced motion correction in cryo-EM single-particle analysis. IUCrJ. 6, 5–17 (2019).

56. J. Zivanov, T. Nakane, S. H. W. Scheres, Estimation of high-order aberrations and anisotropic magnification from cryo-EM data sets in RELION-3.1. IUCrJ. 7, 253–267 (2020).

57. T. Nakane, S. H. W. Scheres, [“Tamir Gonen”, “Brent L. Nannenga”], Eds. (Springer US, New York, NY, 2021; https://doi.org/10.1007/978-1-0716-0966-8_7), vol. 2215 of *Methods in Molecular Biology*, pp. 145–160.

58. B. Webb, A. Sali, Comparative protein structure modeling using MODELLER. Curr Protoc Bioinform. 54: 5.6.1–5.6.37 (2016).

59. F. Madeira, Y. mi Park, J. Lee, N. Buso, T. Gur, N. Madhusoodanan, P. Basutkar, A. R. N. Tivey, S. C. Potter, R. D. Finn, R. Lopez, The EMBL-EBI search and sequence analysis tools APIs in 2019. Nucleic Acids Res. 47, W636–W641 (2019).

60. P. Emsley, B. Lohkamp, W. Scott, K. Cowtan, Features and development of Coot. Acta Crystallogr D Biol Crystallogr. 66, 486–501 (2010).

61. F. Long, R. A. Nicholls, P. Emsley, S. Gražulis, A. Merkys, A. Vaitkus, G. N. Murshudov, AceDRG: a stereochemical description generator for ligands. Acta Crystallogr Sect D Struct Biol. 73, 112–122 (2017).

62. P. V. Afonine, B. K. Poon, R. J. Read, O. V. Sobolev, T. C. Terwilliger, A. Urzhumtsev, P. D. Adams, Real-space refinement in PHENIX for cryo-EM and crystallography. Acta Crystallogr Sect D Struct Biol. 74, 531–544 (2018).

63. C. J. Williams, J. J. Headd, N. W. Moriarty, M. G. Prisant, L. L. Videau, L. N. Deis, V. Verma, D. A. Keedy, B. J. Hintze, V. B. Chen, S. Jain, S. M. Lewis, W. B. Arendall, J. Snoeyink, P. D. Adams, S. C. Lovell, J. S. Richardson, D. C. Richardson, MolProbity: More and better reference data for improved all atom structure validation. Protein Sci. 27, 293–315 (2018).

64. E. F. Pettersen, T. D. Goddard, C. C. Huang, E. C. Meng, G. S. Couch, T. I. Croll, J. H. Morris, T. E. Ferrin, UCSF ChimeraX: Structure visualization for researchers, educators, and developers. Protein Sci. 30, 70–82 (2021).

65. E. Jurrus, D. Engel, K. Star, K. Monson, J. Brandi, L. E. Felberg, D. H. Brookes, L. Wilson, J. Chen, K. Liles, M. Chun, P. Li, D. W. Gohara, T. Dolinsky, R. Konecny, D. R. Koes, J. E. Nielsen, T. Head Gordon, W. Geng, R. Krasny, G. Wei, M. J. Holst, J. A. McCammon, N. A. Baker, Improvements to the APBS biomolecular solvation software suite. Protein Sci. 27, 112–128 (2018).

66. X. Robert, P. Gouet, Deciphering key features in protein structures with the new ENDscript server. Nucleic Acids Res. 42, W320–W324 (2014).

67. A. Goehring, C.-H. Lee, K. H. Wang, J. C. Michel, D. P. Claxton, I. Baconguis, T. Althoff, S. Fischer, K. C. Garcia, E. Gouaux, Screening and large-scale expression of membrane proteins in mammalian cells for structural studies. Nat Protoc. 9, 2574–2585 (2014).

68. S. Nagamori, P. Wiriyasermkul, S. Okuda, N. Kojima, Y. Hari, S. Kiyonaka, Y. Mori, H. Tominaga, R. Ohgaki, Y. Kanai, Structure–activity relations of leucine derivatives reveal critical moieties for cellular uptake and activation of mTORC1-mediated signaling. Amino Acids. 48, 1045–1058 (2016).

69. P. Kongpracha, S. Nagamori, P. Wiriyasermkul, Y. Tanaka, K. Kaneda, S. Okuda, R. Ohgaki, Y. Kanai, Structure-activity relationship of a novel series of inhibitors for cancer type transporter L-type amino acid transporter 1 (LAT1). J Pharmacol Sci. 133, 96–102 (2017).

