## Supplementary Figures 1-13 and Supplementary Tables 1 and 2 for "Ca^2+^-mediated higher-order assembly of b^0,+^AT–rBAT is a key step for system b^0,+^ biogenesis and cystinuria"

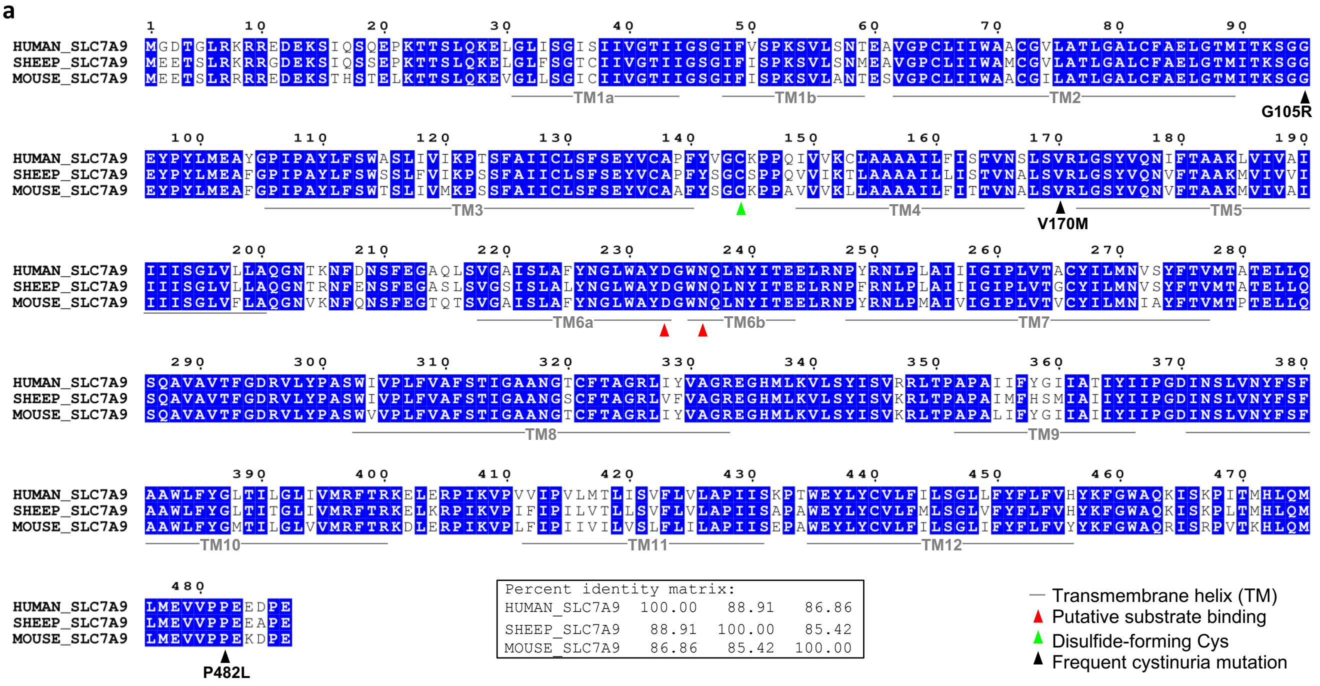


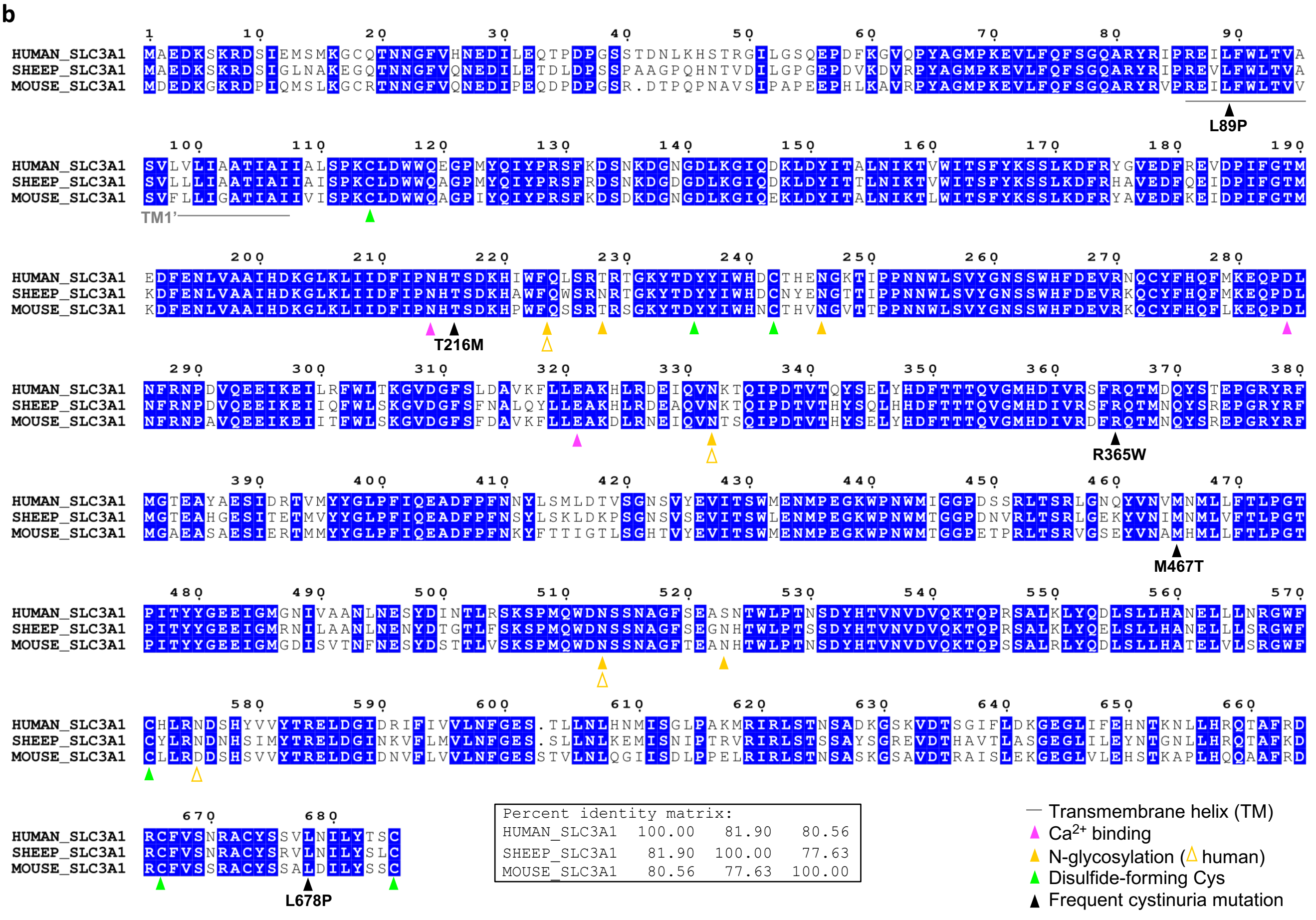


**Figure S1 | Sequence alignments of b$^{0,+}$AT and rBAT**

**a)** Multiple sequence alignment of human (HUMAN_SLC7A9), ovine (SHEEP_SLC7A9), and murine b$^{0,+}$AT (MOUSE_SLC7A9). Alignments were generated by Clustal Omega and formatted by ESPript. Conserved residues are colored blue. The small inset shows the sequence identities (%) between each pair of sequences.
**b)** Multiple sequence alignment of human (HUMAN_SLC3A1), ovine (SHEEP_SLC3A1), and murine rBAT (MOUSE_SLC3A1).

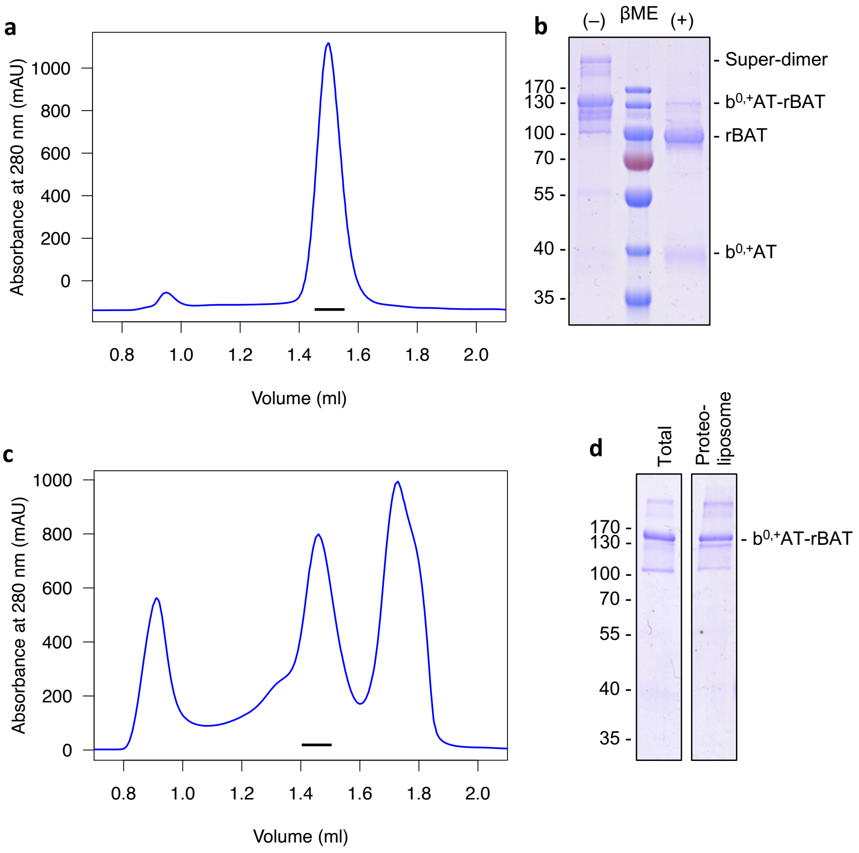


**Figure S2 | Sample preparation of b$^{0,+}$AT–rBAT**

**a)** Size-exclusion chromatography (SEC) profile of purified ovine b$^{0,+}$AT–rBAT complex. The monodisperse peak marked by a black bar was used for cryo-EM and biochemical analyses.

**b)** SDS-PAGE analysis of b$^{0,+}$AT–rBAT. The purified complex was subjected to non-reducing (left) and reducing SDS-PAGE (right).

**c)** SEC profile of nanodisc-reconstituted b$^{0,+}$AT–rBAT. Peak fractions marked by a black bar were used for cryo-EM analysis. The lower molecular-mass peak is excess MSP1E3D1 scaffold protein.

**d)** Reconstitution of b$^{0,+}$AT–rBAT into proteoliposomes. Samples were run on non-reducing SDS-PAGE before and after liposome reconstitution, indicating a reconstitution efficiency of nearly 100%.

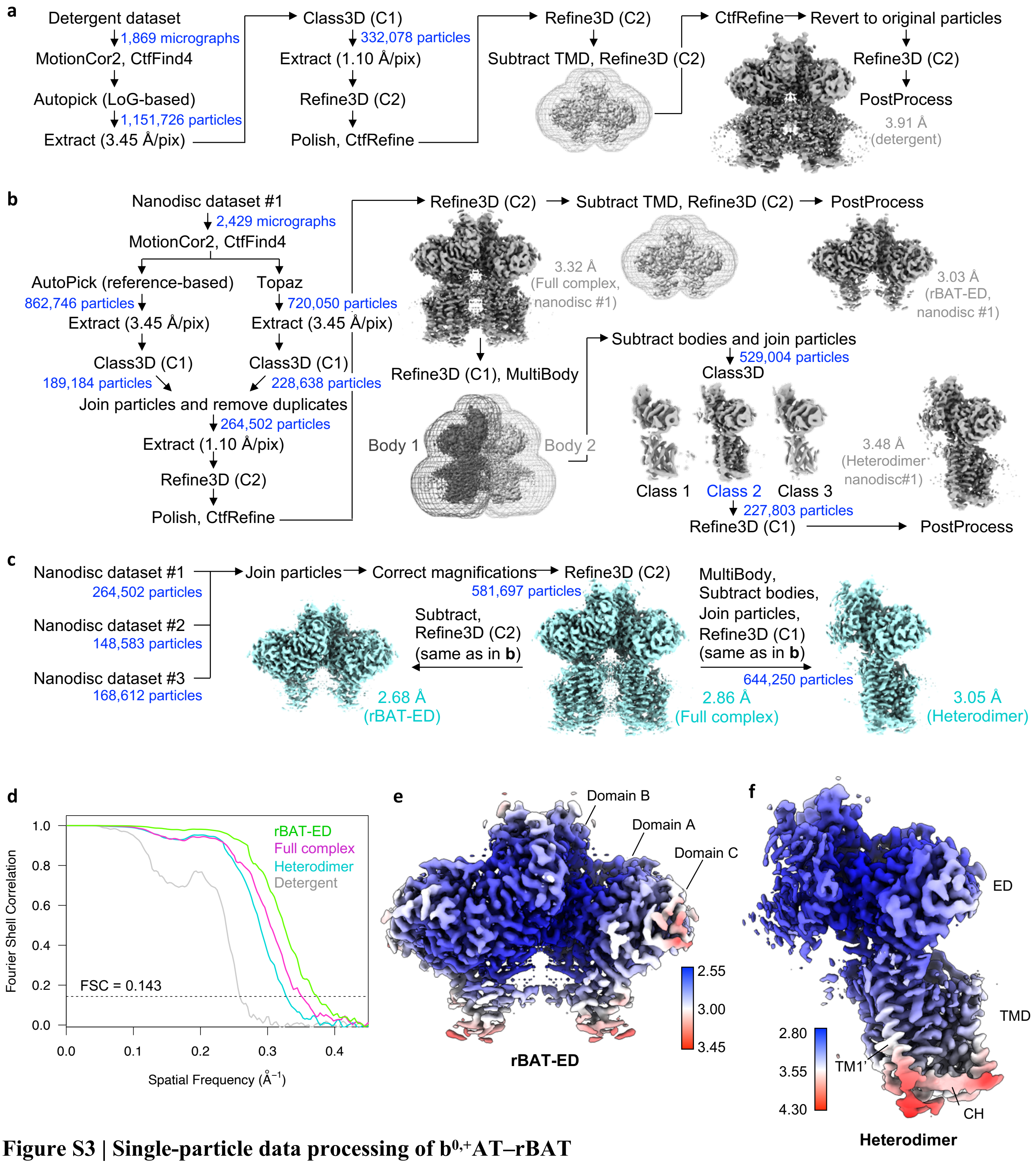


**Figure S3 | Single-particle data processing of b$^{0,+}$AT–rBAT**

**a)** Processing workflow of the detergent dataset.

**b)** Processing workflow of nanodisc dataset #1.

**c)** Merging procedure for nanodisc datasets #1–#3, acquired on three different electron microscopes. Merged data yielded better resolution than individual sets, after correcting for small errors in pixel sizes. Final maps used for model building are colored in cyan. Resolutions for half-map FSC = 0.143 are shown.

**d)** Gold-standard FSC curves calculated for individual maps.

**e)** Local resolution of rBAT-ED dimer.

**f)** Local resolution of heterodimer subcomplex.

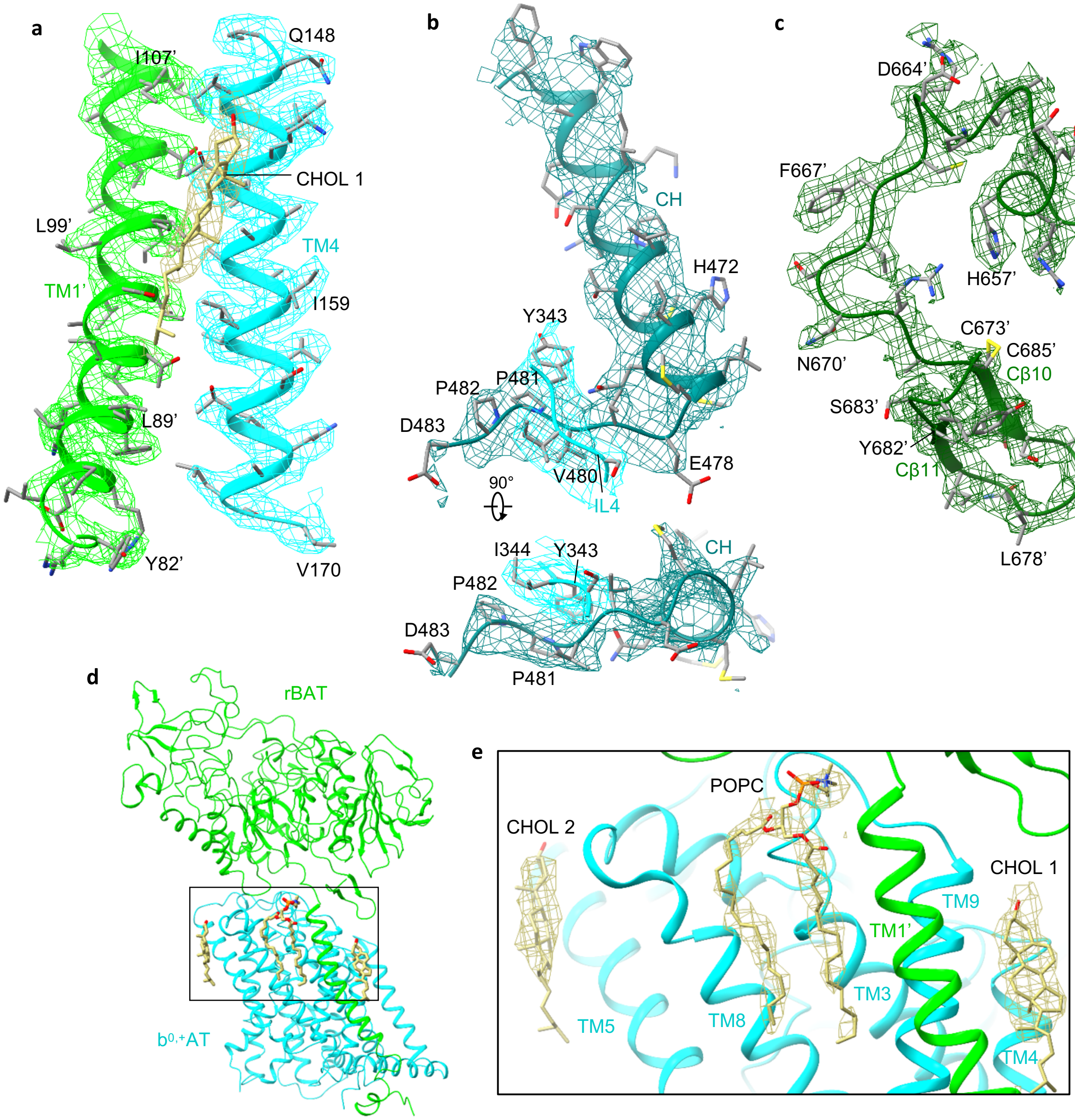


**Figure S4 | Cryo-EM maps in key regions of b$^{0,+}$AT–rBAT**

**a)** TM1'–TM4 interface with a bound cholesterol (CHOL1). Leu89' is highlighted.

**b)** C-terminal helix (CH) of b$^{0,+}$AT. The Val-Pro-Pro motif interacts with intracellular loop 4 (IL4).

**c)** C-terminal peptide of rBAT.

**d)** Three lipids modelled into the density

**d)** Cryo-EM map of the three lipids, derived from the heterodimer map (Fig. S3f).

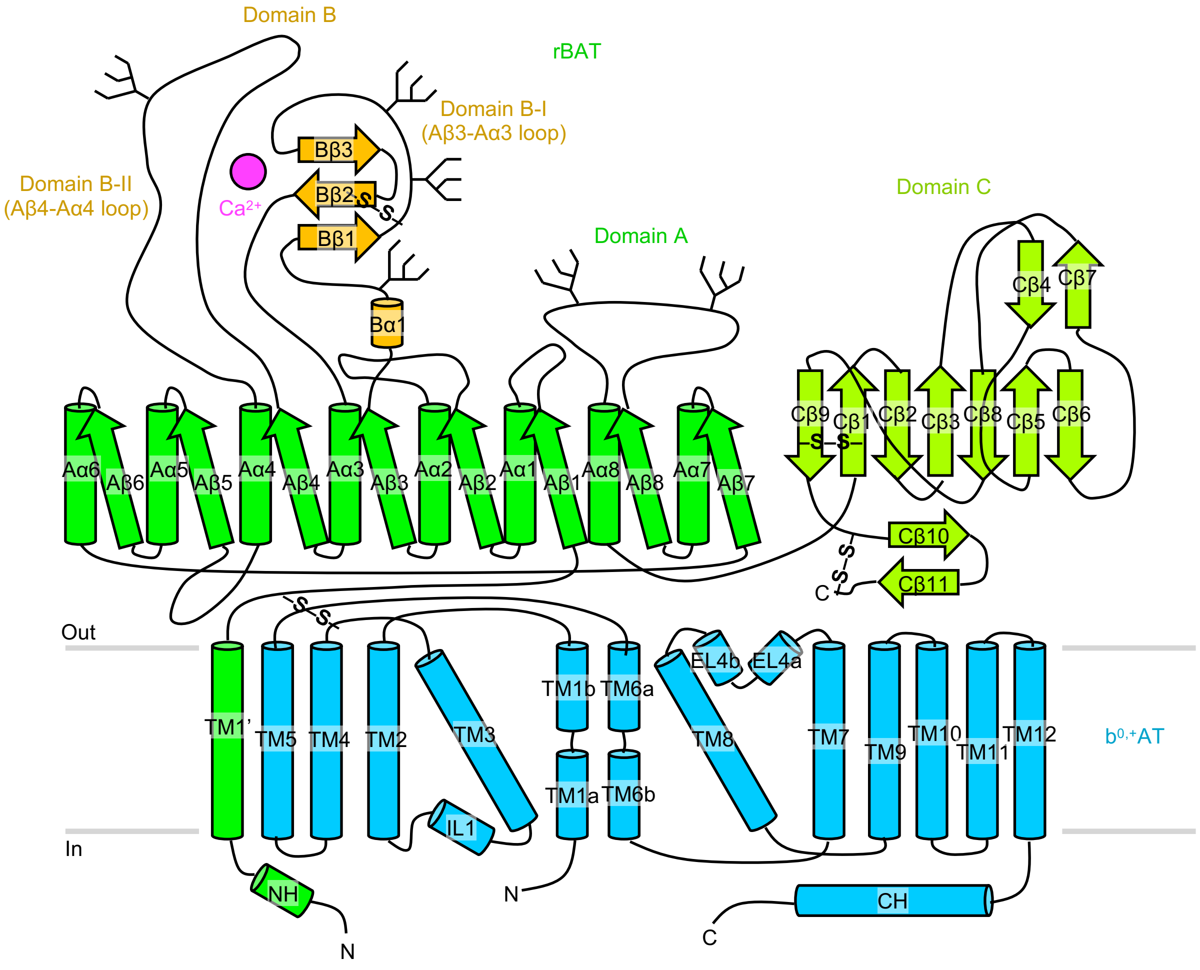


**Figure S5 | Domain architecture of ovine $b^{0,+}$AT–rBAT**

Domain diagram of $b^{0,+}$AT–rBAT, with focus on secondary structure elements, subdomains and post-translational modification (N-linked glycosylation, disulfide bonds and $Ca^{2+}$ binding). rBAT consists of subdomains A, B and C, TM1’, NH and the disordered N-terminal region. Domain B is composed of two loops designated as domain B-I and B-II. $b^{0,+}$AT consists of twelve transmembrane helices (TM1–TM12), two short helices in extracellular loop 4 (EL4a and EL4b), one helix in the intracellular loop 1 (IL1) and the C-terminal helix (CH). The N terminal ~50 residues are disordered in $b^{0,+}$AT.

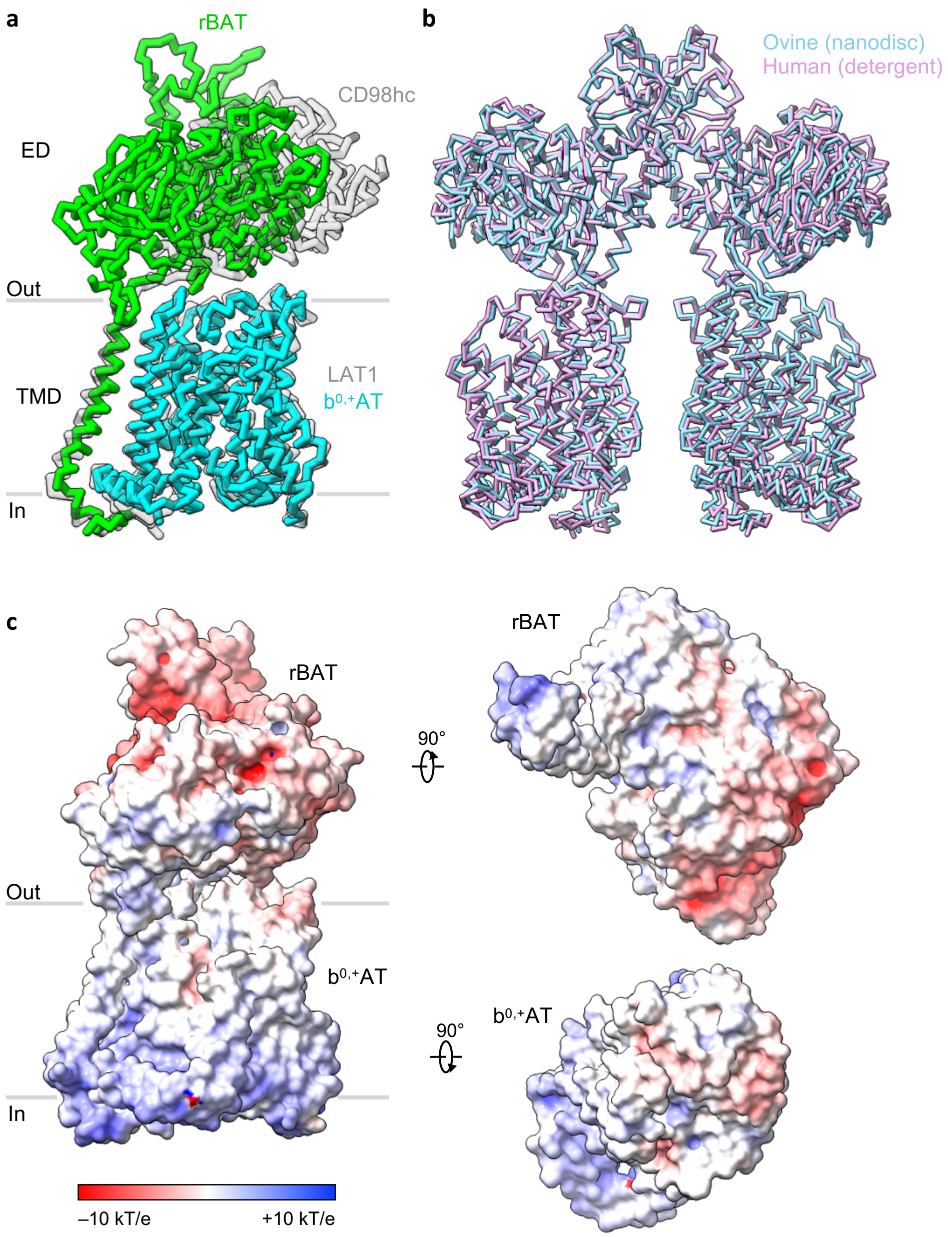


**Figure S6 | Assembly mechanism of b$^{0,+}$AT–rBAT**

**a)** Structural difference between b$^{0,+}$AT–rBAT and LAT1–CD98hc. Structures were superimposed on the TMD. The rBAT ectodomain shows a shift towards TM1' compared to CD98hc, due to the shorter linker between TM1' and ED *(9)*.

**b)** Hetero-tetrameric assembly of ovine and human b$^{0,+}$AT–rBAT (PDB 6LID). Structures were superimposed by the Matchmaker tool in ChimeraX. The two structures are almost identical. The RMSD between 2,152 aligned atom pairs is 1.350 Å.

**c)** Electrostatic potentials calculated for the b$^{0,+}$AT–rBAT heterodimer or separate subunits. Electrostatic potentials were calculated with APBS.

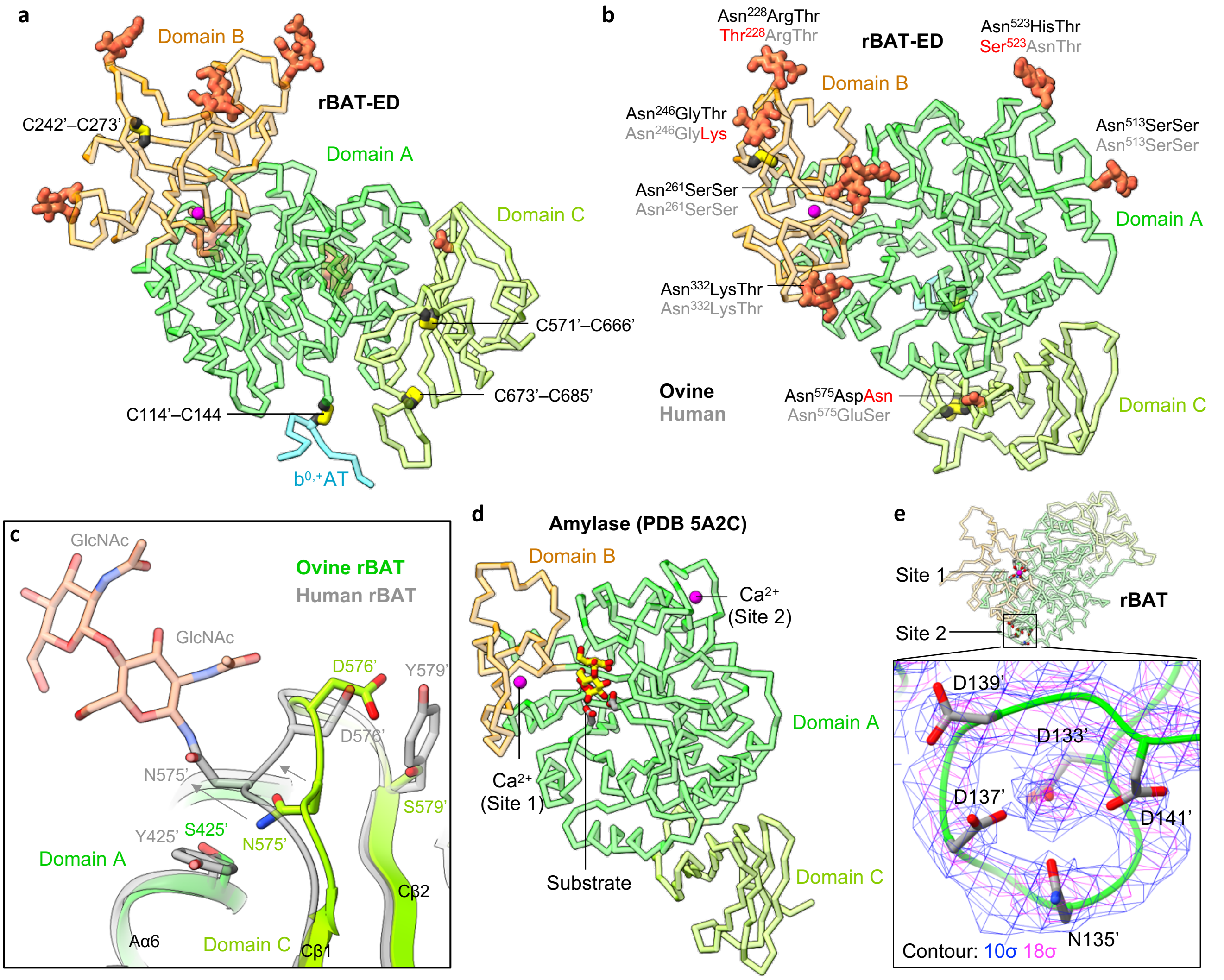


**Figure S7 | Disulfide bonds, N-linked glycans and metal-binding sites of rBAT**

**a)** Disulfide bonds of rBAT. The bond at C114'–C144 is formed across subunits (rBAT and $b^{0,+}$AT), whereas the other bonds are formed within the subunit.

**b)** N-linked glycans of ovine rBAT. The underlying glycosylation motif NX(T/S) is shown for each site. Human rBAT sequences are shown in gray for comparison. Motif-breaking amino acid substitutions are labeled red. Asn575' in ovine rBAT is not glycosylated, whereas the corresponding residue in human is glycosylated.

**c)** Structural differences around Asn575' in domain C. Asn575' of ovine rBAT is not glycosylated and resides within the β-sandwich fold of domain C. In human rBAT, the corresponding residue is glycosylated, protruding towards the extracellular solvent, with acetylglucosamine moieties (GlcNAc) interacting with domain A. The sequence comparison shows that ovine rBAT has small residues around Asn575', namely Ser425' and Ser579', whereas human rBAT has bulky residues Tyr425' and Tyr579', which would result in steric clashes.

**d)** Structure of *Anoxybacillus* α-amylase, which resembles that of the rBAT ectodomain and has high-affinity $Ca^{2+}$ binding sites *(24)* (PDB ID: 5A2C). Site 1 (called Ca1 in the cited literature) is at the domain A–B interface and site 2 (Ca2) is in domain A. Site 1 is close to the substrate, shown as a yellow stick model.

**e)** Weak non-protein density at site 2 of ovine rBAT. The map is contoured at 10 σ and 18 σ. A similar map of site 1 is shown in **Fig. 5a**. Note that this site 2 structurally resembles an EF-hand motif, although it has no strict sequence homology.

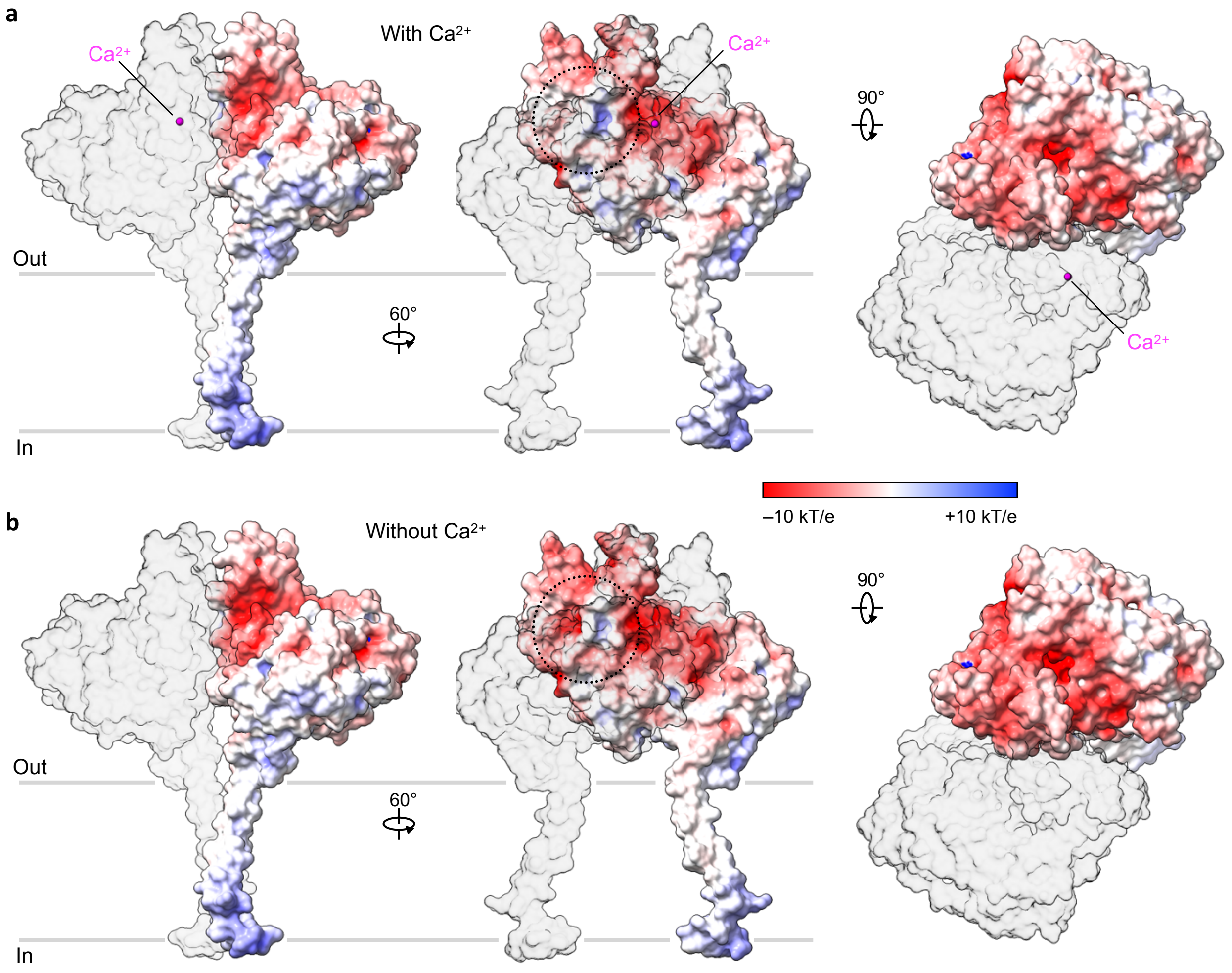


**Figure S8 | Surface electrostatic potential of rBAT**

**a,b)** Electrostatic potential of rBAT with (**a**) and without $Ca^{2+}$ (**b**). The surface patch near the $Ca^{2+}$ binding site shows a strong negative charge in the absence of $Ca^{2+}$ (**a**, middle panel, dotted circle), whereas it is neutralized in the presence of $Ca^{2+}$ (**b** , middle panel, dotted circle). Electrostatic potentials were calculated with APBS.

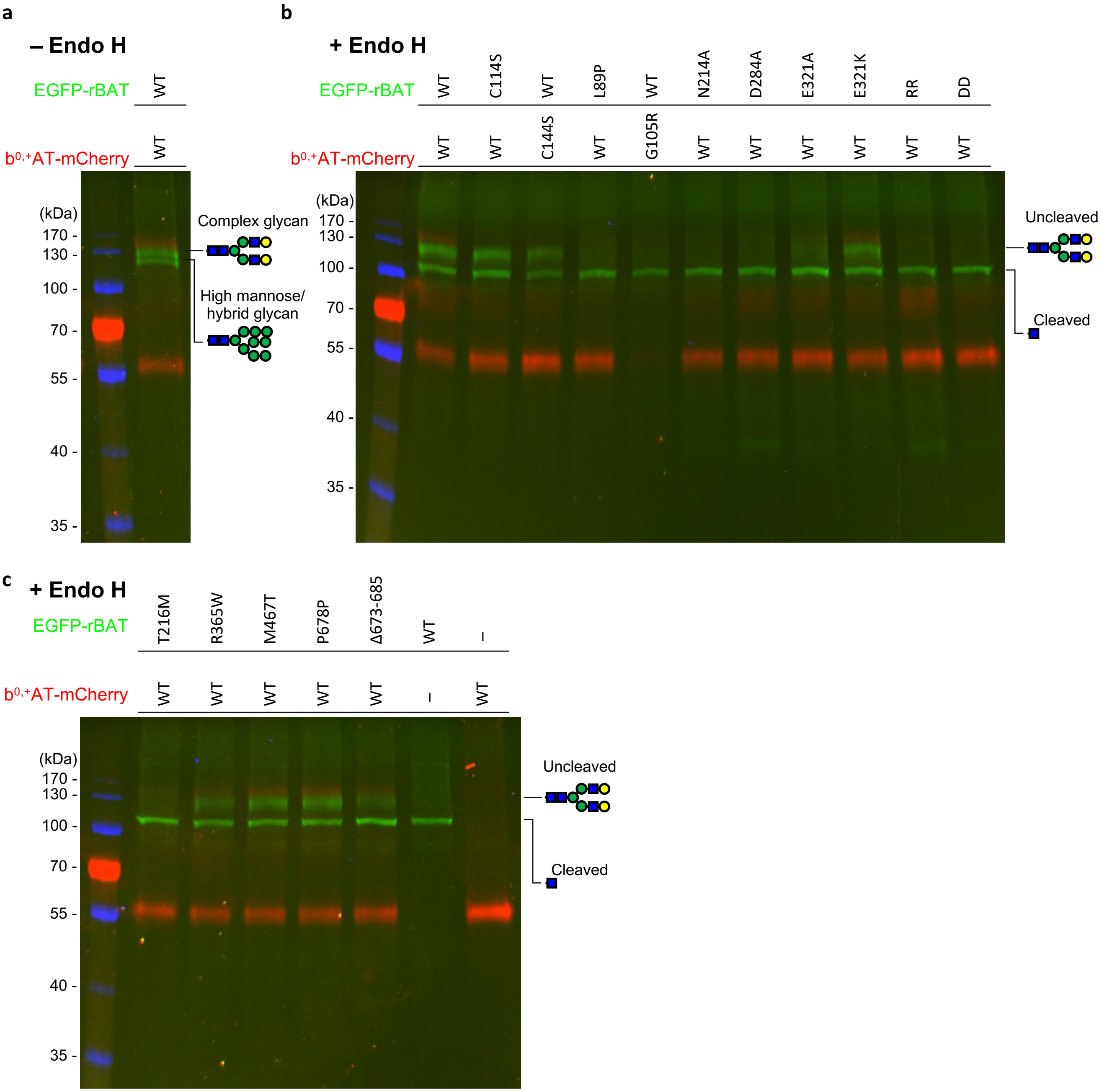


**Figure S9 | Endo H sensitivity assay**

**a)** SDS-PAGE gel of cell lysate expressing EGFP-rBAT and $b^{0,+}$AT-mCherry before Endo H treatment. The image is an overlay of three fluorescence channels for EGFP (green, wavelength = 488 nm, exposure time = 120 sec), mCherry (red, 546 nm, 1440 sec) and the pre-stained molecular weight marker (blue, ex 680 nm, 2 sec). The GFP channel of the same gel is shown in **Fig. 5c**.

**b,c)** SDS-PAGE gels for rBAT and $b^{0,+}$AT, or their mutants, after Endo H treatment. GFP channels of the same gels are shown in **Fig. 5d–f**.

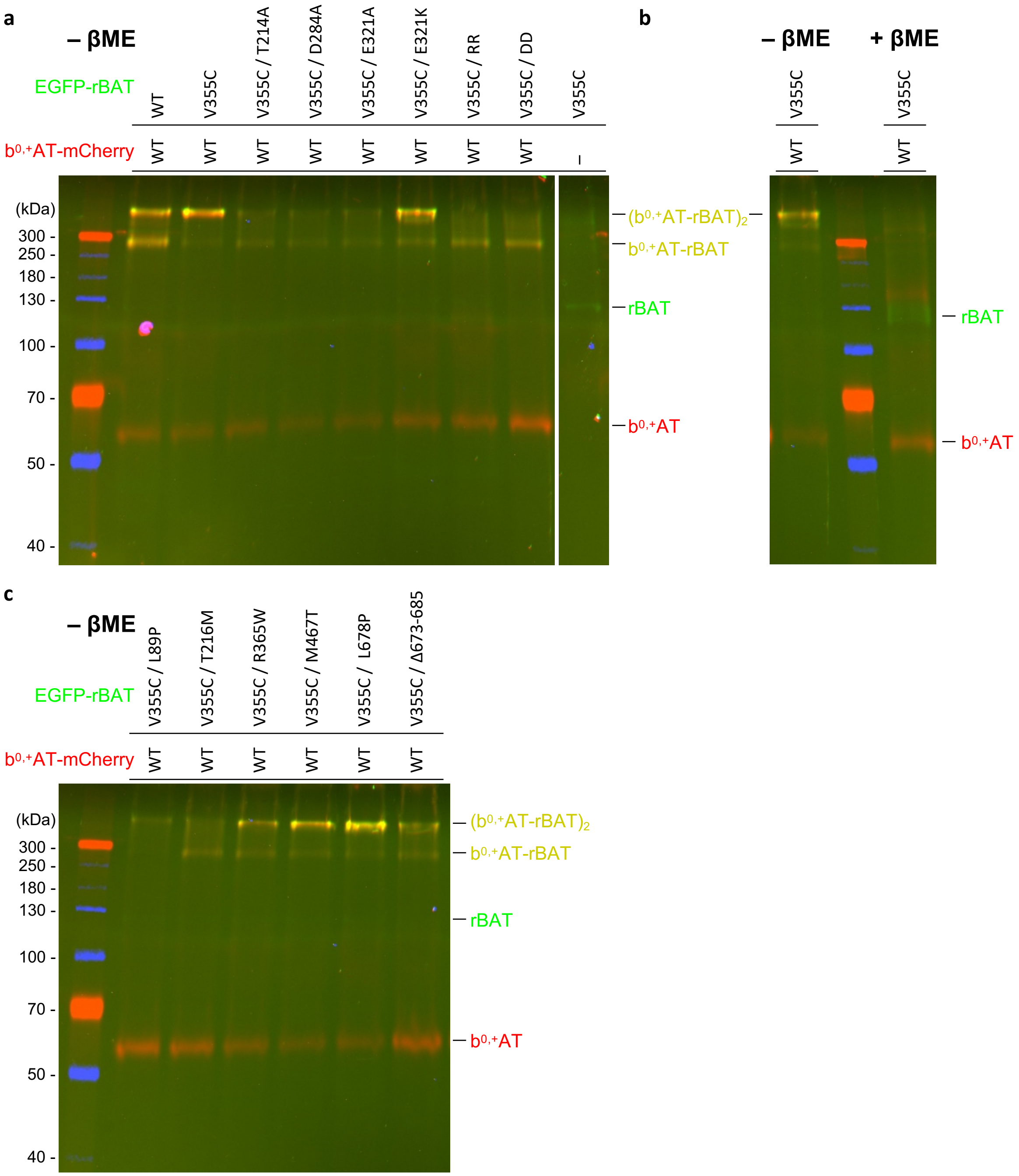


**Figure S10 | Site-specific disulfide cross-linking assay**

**a)** Oxidizing SDS-PAGE for EGFP-rBAT and $b^{0,+}$AT-mCherry, or their mutants, co-expressed in HeLa cells. The three fluorescence channels were imaged as in **Fig. S9a–c**. The yellow bands thus correspond to the heteromeric complexes, of which the upper one is the super-dimeric ($b^{0,+}$AT-rBAT)$_2$ complex, whereas the lower band is the $b^{0,+}$AT-rBAT subcomplex. A cropped image of the same gel is shown in **Fig. 5h,j**.

**b)** Cross-linking validation. Upon addition of β-mercaptoethanol, the yellow band dissociates into two bands, each representing EGFP-rBAT (green) or $b^{0,+}$AT-mCherry (red), confirming specific disulfide cross-linking. A cropped image of the same gel is shown in **Fig. 5k**.

**c)** Oxidizing SDS-PAGE for cystinuria mutants. A cropped image of the same gel is shown in **Fig. 5i**.

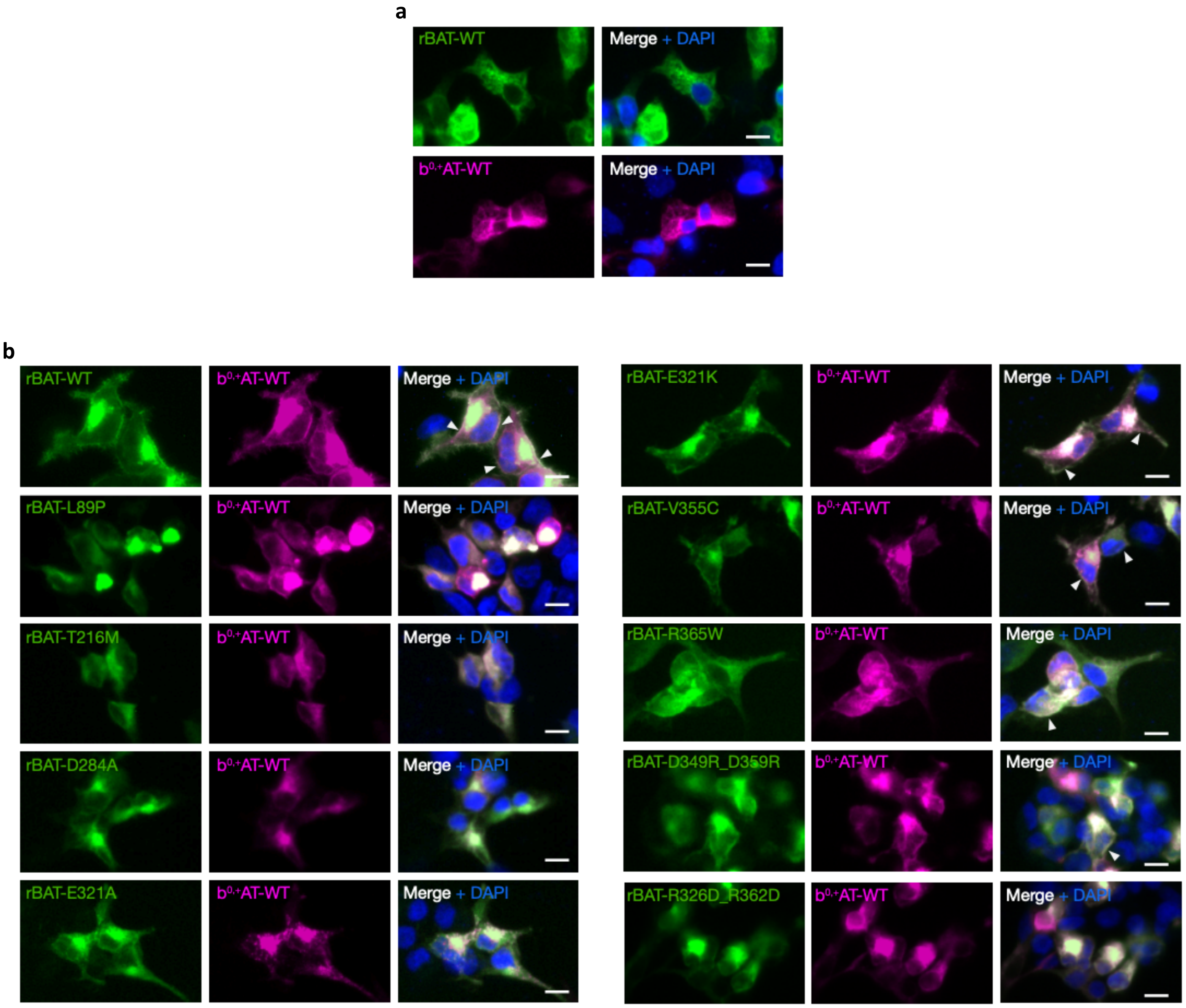


**Figure S11 | Fluorescent cell imaging**

**a)** Fluorescence imaging of HeLa cells individually expressing rBAT-WT (EGFP-tagged) or $b^{0,+}$AT-WT (mCherry-tagged) as negative controls.

**b)** Fluorescence imaging of HeLa cells co-expressing rBAT and $b^{0,+}$AT, either as wild-type or mutants. Cell-surface co-localization of rBAT and $b^{0,+}$AT is marked by white arrows. Trafficking-deficient mutants show reduced cell-surface co-localization. High accumulation of rBAT and $b^{0,+}$AT inside of cells are due to over-expression. Scale bars = 10 µm.

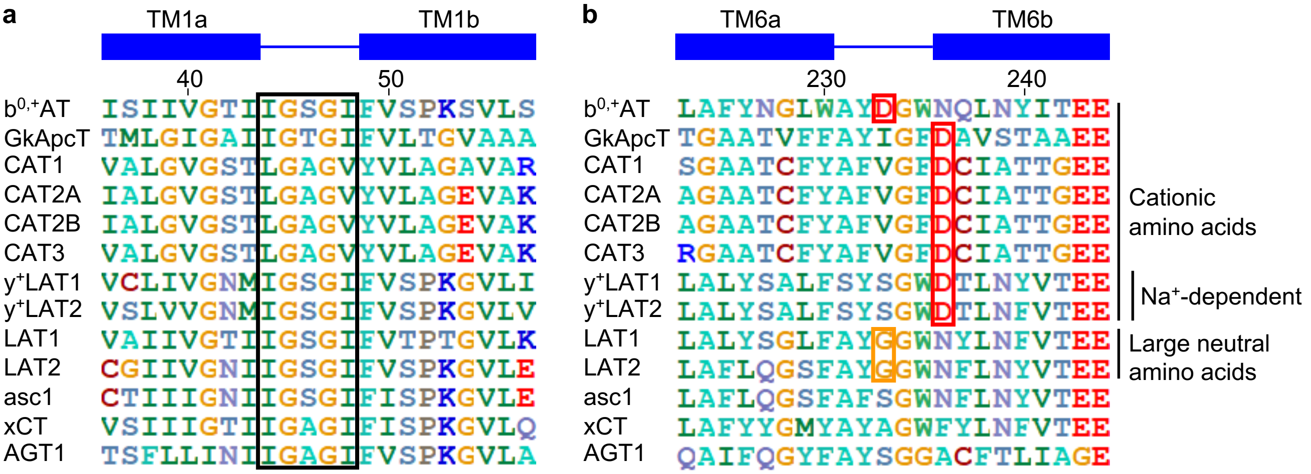


**Figure S12 | Sequence features of TM1 and TM6 in SLC7 transporters**

**a)** Sequence alignment of TM1 in human SLC7 (SLC7A1–11, 13) and GkApcT. The unwound region between TM1a and TM1b, highlighted by a rectangle, is well-conserved across all SLC7 members.

**b)** Sequence alignment of TM6 in human SLC7 (SLC7A1–11, 13) and GkApcT. Acidic residues implicated in cation recognition are outlined in red. Gly residues important in system L are are outlined in orange.

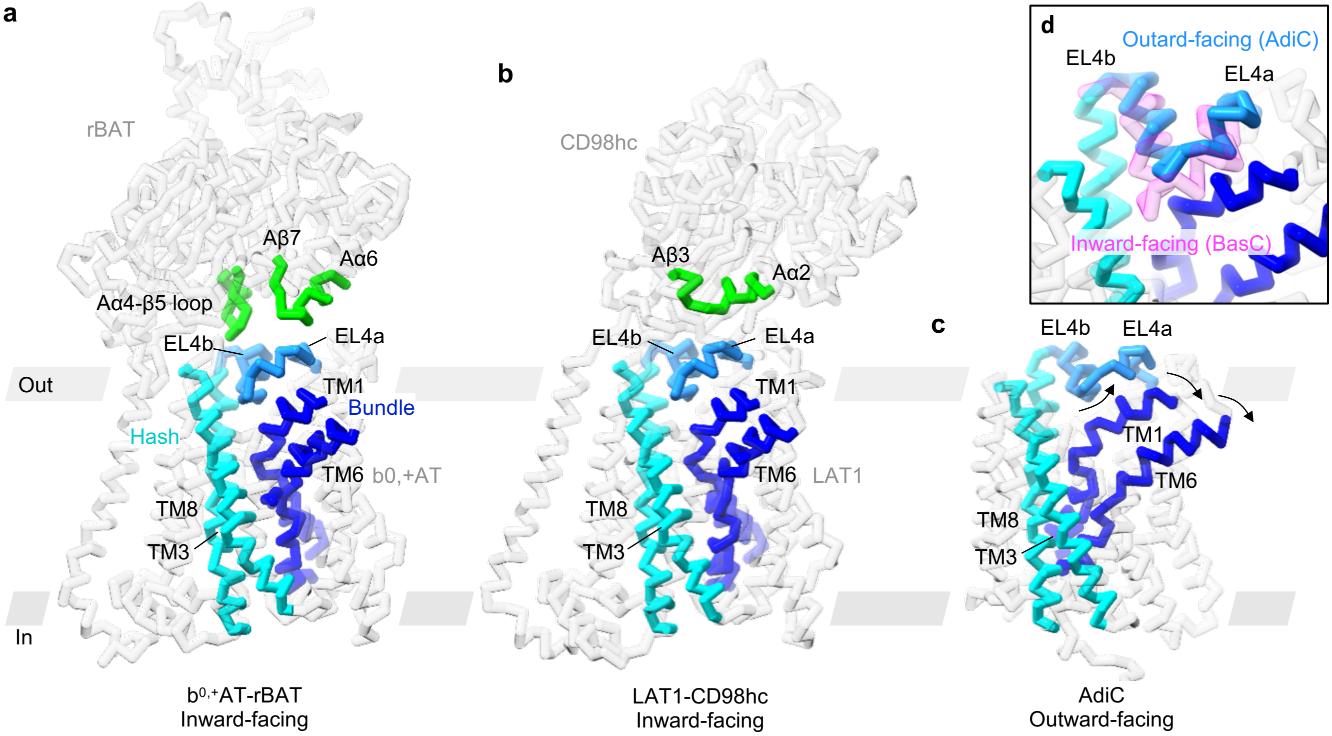


**Figure S13 | Possible mechanisms of SLC7 regulation by SLC3**

**a)** Structure of $b^{0,+}$AT–rBAT, highlighting structural elements important for substrate translocation. Extracellular halves of TM1 and TM6 are close to TM3 and TM8, closing the extracellular gate. EL4a and EL4b act as a lid, stabilized by two loops of rBAT.

**b)** Inward-facing structure of LAT1–CD98hc. As in $b^{0,+}$AT, EL4a and EL4b of LAT1 form a lid above the extracellular gate, while interacting with a different CD98hc loop.

**c)** Structure of AdiC in the outward-facing conformation. Swinging movements of TM1, TM6, EL4a and EL4b define the opening of the extracellular gate and are depicted by arrows. Along with this movement, the extracellular halves of TM1 and TM6 dissociate from TM3 and TM8 to open the extracellular gate. EL4a and EL4b show concerted movements to widen the substrate pathway.

**d)** Close-up view of EL4a and EL4b in two bacterial SLC7 homologues. Comparison of the outward-facing structure of AdiC (cyan) and the inward-facing structure of BasC (pink) shows that EL4a and EL4b undergo sliding movements to enable a wide opening of the extracellular gate.

**Table S1 | Cryo-EM data collection**

| | Detergent | Nanodisc #1 | Nanodisc #2 | Nanodisc #3 |
|---|---|---|---|---|
| **Data collection** | | | | |
| Microscope | Titan Krios 2 | Titan Krios 2 | Titan Krios 1 | Titan Krios 2 |
| Camera | K3 | K3 | K3 | Falcon III |
| Magnification | 105,000 | 105,000 | 105,000 | 96,000 |
| Voltage (kV) | 300 | 300 | 300 | 300 |
| Electron exposure ($e^{-}/Å^2$) | 50.0 | 50.0 | 40.0 | 40.0 |
| Defocus range (μm) | −1.4 to −2.5 | −1.2 to −2.2 | −1.0 to −2.0 | −1.4 to −2.0 |
| Calibrated pixel size (Å) | 0.837 | 0.837 | 0.831 | 0.833 |
| Initial particle images (no.) | | | | |
| RELION / Topaz | 1,151,726 / – | 862,746 / 720,050 | 456,825 / 579,851 | 855,952 / – |

**Table S2 | Data processing, model building and validation statistics**

| | Detergent | Nanodisc #1 – #3 merged | | |
|---|---|---|---|---|
| | | Full complex | rBAT ectodomain | Heterodimer |
| **Data processing** | | | | |
| Final particle images (no.) | 332,078 | 581,697 | 581,697 | 644,250 |
| Final pixel size (Å) | 1.09856 | 1.09856 | 1.09856 | 1.09856 |
| Symmetry imposed | C2 | C2 | C2 | C1 |
| Map resolution (Å) | | | | |
| Half map FSC = 0.143 | 3.91 | 2.86 | 2.68 | 3.05 |
| Map sharpening *B* factor ($Å^2$) | –111.3 | –63.1 | –76.9 | –85.1 |
| Local resolution range (Å) | | | 2.55–3.45 | 2.80–4.30 |
| **Refinement** | | | | |
| Initial model (PDB codes) | | | 1UOK, 6AAV, 6IRS | 1UOK, 6AAV, 6IRS |
| Refinement resolution (Å) | | | 2.60 | 3.00 |
| Model resolution (Å) | | | | |
| Map-model FSC = 0.5 | | | 2.62 | 2.95 |
| Model composition | | | | |
| Non-hydrogen atoms | | | 4,821[a] | 8,817 |
| Protein residues | | | 575 | 1,078 |
| No. ligands | | | GlcNAc: 8<br>$Ca^{2+}$: 1 | GlcNAc: 8<br>$Ca^{2+}$: 1<br>POPC: 1<br>Cholesterol: 2 |
| Average *B* factors ($Å^2$) | | | | |
| Protein | | | 50.24 | 72.85 |
| Ligand | | | 86.60 | 100.92 |
| R.m.s. deviation | | | | |
| Bond lengths (Å) | | | 0.005 | 0.006 |
| Bond angles (°) | | | 0.672 | 0.718 |
| **Validation** | | | | |
| MolProbity score | | | 2.25 | 2.46 |
| Clashscore | | | 5.36[b] | 7.61 |
| Rotamer outliers (%) | | | 6.53 | 8.17 |
| Cβ outliers (%) | | | 0.00 | 0.00 |
| CaBLAM outliers (%) | | | 3.68 | 2.62 |
| Ramachandran plot | | | | |
| Favored (%) | | | 95.11 | 94.88 |
| Allowed (%) | | | 4.71 | 5.03 |
| Outliers (%) | | | 0.17 | 0.09 |

[a] for one asymmetric unit.
[b] Clashscore is calculated for a C2-expanded homodimer to take homomeric interfaces into account.
